# Developing a fluorescent derivative of GagPol as a tool for live-cell imaging of HIV-1 assembly

**DOI:** 10.64898/2026.02.09.704847

**Authors:** Irene Gialdini, Virgile Rat, Maria Anders-Össwein, Barbara Müller, Don C. Lamb

**Author notes:** These authors contributed equally to this work (Irene Gialdini, Virgile Rat).

## Abstract

The HIV-1 assembly process is driven by the structural polyprotein Gag, which forms small cytosolic oligomers that traffic to the plasma membrane. While Gag alone is sufficient to drive particle formation, the incorporation of GagPol, a polyprotein comprising Gag and viral enzymes, is essential for productive infection. Maintenance of the proper Gag:GagPol ratio is crucial for virion infectivity. Yet, the mechanisms regulating their cytosolic interactions remain incompletely understood, in part because most studies rely on ensemble biochemical assays that hide cell-to-cell heterogeneity and lack spatial and temporal resolution. To overcome these limitations and investigate the dynamics of GagPol at the molecule level, we systematically tested different approaches to fluorescently label it. We successfully produced two functional fluorescent GagPol variants: one single-labeled GagPol, and a second double-labeled variant with a fluorescent protein added within Gag, that allows concurrent visualization of Gag and GagPol. These labeled versions, in combination with the use of raster image (cross-) correlation spectroscopy, enabled the quantification of Gag and GagPol relative concentrations and intermolecular interactions at the single-cell level. Overall, these variants set the stage for in-depth investigations of GagPol during HIV assembly providing insights into cytoplasmic trafficking, particle assembly, and the kinetics of these processes.

**Importance:** The participation of Gag and GagPol in HIV-1 assembly is central to the generation of infectious virions. While the structural role of Gag and the enzymatic functions of GagPol are well described, little is known regarding the mechanisms underlying their cytosolic interactions, spatial organization, and relative incorporation into assembling particles. Here, we successfully developed two stable fluorescently labeled variants of GagPol; a single labeled variant with a label in the Pol domain and double-labeled construct in which the Gag domain is also tagged. Using these constructs, we can quantitatively investigate Gag and GagPol interactions on the single-cell level. Hence, they provide a framework for a comprehensive study of the oligomerization properties and trafficking of these polyproteins at different stages of HIV-1 assembly, including membrane recruitment and incorporation into nascent virions.

## Introduction

Human Immunodeficiency virus type 1 (HIV-1) is the etiological agent of the acquired immunodeficiency syndrome (AIDS). With an estimated total of 40.8 million people living with HIV and ∼630,000 deaths from HIV-related causes in 2024, the virus still presents a major health concern (1). HIV particles are heterogeneous in size, with an average diameter of roughly 145 nm (2). The lipid envelope carrying the envelope glycoprotein Env surrounds a protein shell consisting of the domains of the major structural polyprotein Gag: MA (matrix), CA (capsid), NC (nucleocapsid) and p6. Other main components of the virion are the enzymes PR (protease), RT (reverse transcriptase) and IN (integrase), encoded as parts of a 160 kDa GagPol polyprotein, and two copies of a positive-sense, single-stranded genomic RNA (gRNA).

Virion assembly at the plasma membrane is orchestrated by the 55 kDa Gag polyprotein. The subdomains of Gag have distinct functions in virion assembly and release: MA mediates Gag plasma membrane targeting and membrane association, CA contributes protein-protein interaction interfaces important for particle assembly, NC binds to the genomic RNA (gRNA) and p6 recruits components of the cellular sorting complexes endosomal required for transport (ESCRT) that mediate budding of the virion from the plasma membrane. Roughly 2-3000 Gag molecules are the main constituent of the virion; the protein also mediates incorporation of GagPol and the gRNA into the nascent particle (3–6). HIV becomes infectious upon proteolytic cleavage of Gag and GagPol into the mature subunits by the viral PR concomitant with budding (3–6), accompanied by morphological rearrangement into the mature virion.

While Gag alone can assemble into virus-like particles, the incorporation of GagPol comprising the viral enzymes in appropriate amounts is essential for virion infectivity. In virus producing cells, Gag-Pol is synthesized via translational frameshift. The 5’ part of the *gag* open reading frame (ORF) overlaps with the *pol* ORF with a shift of -1 bp. During translation, a conserved “slippery sequence” (UUUUUUA) in coordination with an adjacent stem-loop structure within the NC encoding region of *gag* induces ribosomal staggering, promoting a frameshift to the *pol* ORF (7, 8). This frameshift is reported to occur with a frequency of ∼5% of translation events. The frameshift signals are highly conserved and subject to selection for viral fitness and artificially increasing the proportion of Gag-Pol to Gag in expressing cells impairs virion production and infectivity (8–10).

Not only the stoichiometry between Gag and GagPol, but also the dimerization of GagPol in the assembly process needs to be tightly regulated. The PR domain of GagPol is a monomer that needs to dimerize to become activated. Premature or impaired activation of PR affects assembly and maturation and renders the virus non-infectious as reviewed by (11). Although precise control of Gag-GagPol and GagPol-GagPol interactions is crucial for the formation of infectious HIV-1, the mechanisms of this regulation are currently incompletely understood. The nucleation of GagPol dimerization during assembly, the regulation of optimal Gag:GagPol stoichiometry, as well as the role of the viral gRNA in these processes, are still under investigation. The question of whether protein stoichiometry in the virion is achieved solely by the ratio of Gag to GagPol determined by frameshifting efficiency has been addressed using a bulk biochemical approach (10). An immunoblot-based estimate of the relative amounts of Gag and GagPol in virus producing cells and pelleted particles suggested a ∼3-fold enrichment of GagPol in virus preparations compared to cell lysates. This indicates that additional mechanisms beyond stochastic incorporation are involved in regulating GagPol packaging.

The ratio of Gag to GagPol has never been quantified on the single-cell level and only recently been investigated in the context of mature particles (9, 10). These previous studies employed bulk biochemical methods, such as immunoblot, which provide averaged ensemble data that do not consider cell-to-cell heterogeneity. Furthermore, biochemical approaches lack temporal and spatial resolution. Obtaining more detailed quantitative insights into GagPol dimerization, assembly dynamics and incorporation at the single cell and single virus level requires alternative approaches. Fluorescent labeling of HIV-1 Gag and GagPol in the viral context, retaining the original expression ratio and assembly functionality would enable a quantitative spatio-temporal description of HIV-1 assembly using advanced microscopy methods. However, labeling GagPol is challenging. GagPol dimerization and incorporation is driven by protein-protein and protein-RNA interactions involving multiple regions within the polyprotein (12–16). Insertions, or even minor modifications may not only affect assembly properties, but also the stability of the protein (17).

In this work, we systematically tested different approaches and generated a labeled variant that enabled us to visualize both Gag and GagPol expressed in the viral context by fluorescence microscopy. While insertions between PR and RT or RT and IN led to the production of aberrant GagPol products, the substitution of the IN domain with a fluorescent protein did not interfere with overall integrity of GagPol and virion formation. Combining this modification with the insertion of a second fluorophore between MA and CA, which has been previously used to label Gag with minimal perturbation to its functionality (6, 18–21), enables the study of GagPol dynamics and interactions between Gag and GagPol at the single cell level.

## Results

To visualize cytosolic GagPol, a fluorescent protein needs to be introduced into the Pol coding region. Since labeling can potentially impact protein folding and stability and induce proteolytic degradation, we systematically examined multiple sites in the Pol coding region within the context of the HIV-1 subgenomic plasmid pCHIV (Figure 1) (20). pCHIV encodes all HIV-1 proteins with the exception of Nef and does not comprise the long terminal repeat regions essential for virus replication. To study virion assembly independent of maturation, all variants used carried a point mutation of the PR active site (D25N). Each labeling position was investigated using fluorescence imaging of transfected cells expressing labeled GagPol, and immunoblots of virus-like particles (VLPs) and cell lysates.

**Figure 1:**
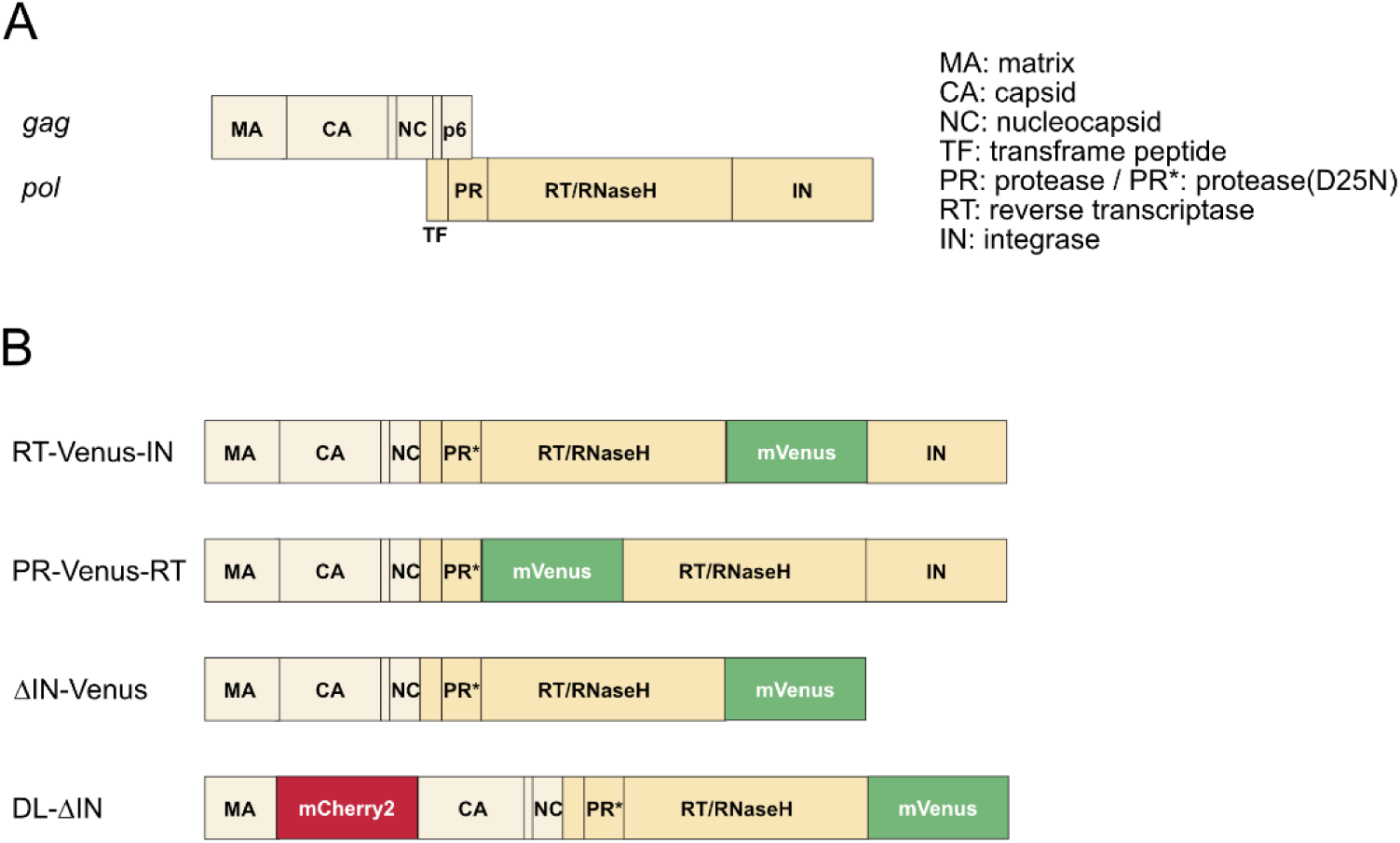
Graphic representation of HIV-1 GagPol domains and labeling positions. A) Wild-type Gag and GagPol domains. B) GagPol derivatives generated in this study.

The diffusion of cytosolic GagPol was assessed via Raster Image Correlation Spectroscopy (RICS) (22) performed in live cells 16-18 h after transfection with a GagPol-labeled construct. RICS is a fluorescence fluctuation method that provides quantitative information regarding mobility and concentration of labeled molecules as illustrated in Figure 2A (for more details on the method, see (23)). A 12 × 12 µm region of interest situated in the cytoplasm was selected and 150 frames were acquired with raster scanning. The raster scanning embeds dynamic information into a single image, which is extracted using a spatial autocorrelation analysis. The spatial autocorrelation function (SACF) is calculated per frame and all SACFs from the measurement are then averaged together. By fitting the average SACF, the diffusion coefficient (*D*) and concentration (*c*) of labeled GagPol are retrieved. Since RICS can be heavily affected by signal heterogeneities such as bright aggregates, we selected cells in early stages of virion production that did not yet exhibit VLPs formation at the plasma membrane.

**Figure 2:**
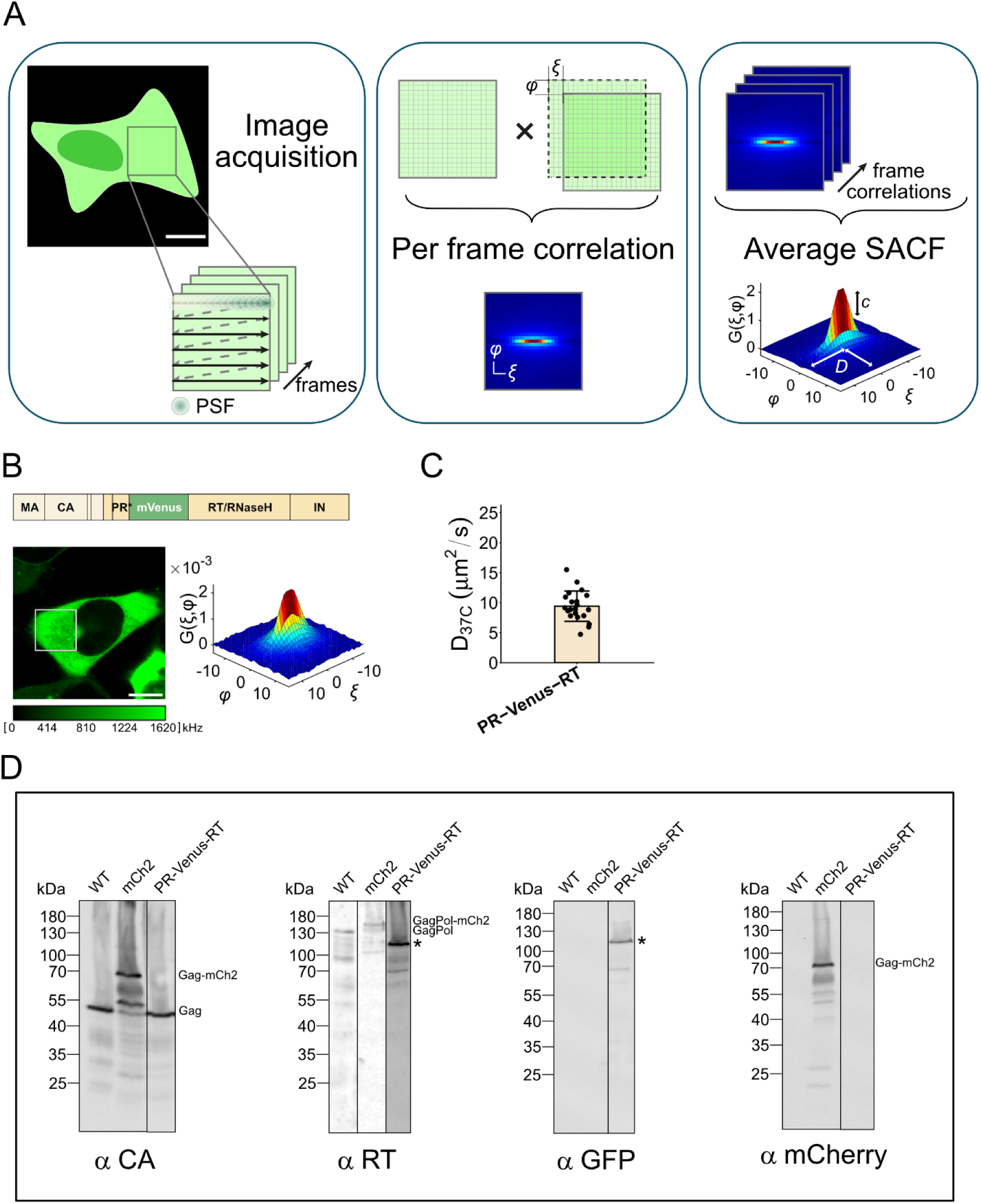
mVenus insertion between PR and RT domains of GagPol. **A)** Schematic representation of the RICS workflow. Left panel: after selecting a 12 × 12 µm region of interest within the cytoplasm of the cell, 150 frames were acquired by raster scanning, as indicated by the arrows. Scanning of the point spread function (PSF) of the laser is indicated. Central panel: Each frame is individually correlated. The correlation is calculated between the acquired frame and a copy of it, shifted it by pixel lags (ξ, ψ). This process is repeated for the desired combinations of (ξ, ψ). Right panel: the correlation functions obtained from each individual frame are averaged, generating the average spatial autocorrelation function (SACF). **B)** Left: Representative image of HeLa cells transfected with the PR-Venus-RT construct. RICS was performed by imaging the ROI highlighted by the white square and the SACF is reported on the right. Scale bar 10 µm. Right: SACF. **C)** Diffusion coefficient (mean ± standard deviation) of GagPol constructs of panel A. Results from individual cells (n = 21), pooled from two biological replicates, are shown as individual data points. **D)** Immunoblot analysis of GagPol derivative PR-Venus-RT in cell lysates. Cells were transfected with a plasmid expressing wild-type or tagged GagPol in the context of pCHIV (20). At 48h post transfection, cell lysates were harvested, proteins were separated by SDS-PAGE and transferred to a nitrocellulose membrane. Gag, GagPol and derivatives were detected by ECL immunoblot using the indicated antisera. Positions of the molecular mass marker are indicated (in kDa). The * indicates the aberrant GagPol product. Individual immunoblots are outlined by a black border.

### Impact of the fluorescent protein inserted at the PR-RT junction

First, we placed mVenus, preceded by a flexible dipeptide linker (Ser-Gly), at the PR-RT junction, as previously described by Takagi et al (24). This monomeric fluorophore is known for its fast maturation rate and high brightness (25, 26) and was previously utilized to label Gag (27). HeLa cells expressing PR-Venus-RT imaged 18h post-transfection displayed a homogeneous cytoplasmic fluorescence signal with minimal fluorescence observed in the cell nucleus (Figure 2B, left panel), in good agreement with Takagi et al. RICS experiments performed on cytoplasmatic ROIs revealed a *D* = 9.4 ± 2.5 µm^2^/s (Figure 2C, Supplementary Table S1), which is faster than expected, based on the diffusion of mCherry2-labeled Gag (*D* = 2.1 ± 1.5 µm^2^/s, expressed from the pCHIV.mCherry2 variant, Supplementary Figure S1 and Supplementary Table S2).

The expression of the tagged GagPol protein was also analyzed by immunoblot of cell lysates. As positive control, we included lysates from cells transfected with the previously characterized pCHIV.mCherry2. In this variant, both Gag and GagPol carry a fluorophore moiety between the MA and CA domains of Gag resulting in a molecular weight similar to the PR-Venus-RT variant. Immunoblots of cell lysates from PR-Venus-RT expressing cells displayed wild-type like expression of Gag (Figure 2D). However, while a faint band corresponding with the full-length constructs was detected, the dominant product had a molecular mass significantly lower than that of GagPol expressed from pCHIV.mCherry2. This product was detected with both anti-RT and an anti-GFP antibodies, suggesting the expression of an aberrant polyprotein, consistent with its high mobility determined in RICS experiments. Thus, we considered this variant unsuitable for our purposes.

Since GagPol PR-RT demonstrated the expected cytosolic fluorescence localization, we tried to rescue the aberrant protein expression by changing the peptide linkers flanking the fluorophore (Supplementary Figure S2). To this end, we extended the Serine-Glycine dipeptide linker of the PR-Venus-RT variant by inserting either an additional Ser-Gly repeat, maintaining the linker flexibility, or by adding a four Proline repeat, generating a short rigid linker upstream of mVenus. We also replaced mVenus with an eCFP moiety flanked by flexible linkers (the same extended Serine-Glycine linker upstream and a Glycine-Serine-Glycine linker downstream). All these variants showed a cytosolic phenotype (Supplementary Figure S2A) and a *D* ranging from 7 to 10 µm^2^/s (Supplementary Figure S2B and Supplementary Table S1). Likewise, the immunoblots revealed that what we were observing was not full length GagPol but an aberrant derivative, similar to the initial version of PR-Venus-RT (Supplementary Figure S2C). Thus, we concluded that the insertion of a heterologous protein between PR and RT compromised the integrity of GagPol.

### Inserting the fluorescent protein flanking the IN domain results in aberrant GagPol expression

After excluding the PR-RT junction as a potential labeling site, we placed mVenus upstream of IN (construct RT-Venus-IN) (Figure 3A, upper panel). The introduction of various fluorescent proteins at this position has been reported to be relatively well tolerated with respect to virion infectivity (28, 29). Unexpectedly, we observed both cytosolic and nuclear fluorescence, suggesting at least partial instability of the polyprotein. Indeed, large polyproteins such as Gag and GagPol are expected to localize solely in the cytosol (24, 27, 30, 31). RICS analyses performed on cytoplasmatic ROIs (Figure 3A, right panel) supported premature cleavage of the RT-Venus-IN constructs, as an average diffusion coefficient of *D* = 8 ± 4 µm^2^/s was extracted (Figure 3B and Supplementary Table S1). Such diffusion is faster than expected for the full-length GagPol but too slow to be attributed to mVenus alone, whose diffusion coefficient was *D* = 36 ± 14 µm^2^/s (Supplementary Figure S1, Supplementary Table S1). Hence, it could represent degradation products of GagPol.mVenus comprising the fluorophore. This result was confirmed by immunoblot analyses of cell lysates from RT-Venus-IN expressing cells, which displayed a dominant product of ∼55 kDa containing mVenus (Figure 3C, anti-GFP blot). Therefore, the RT-Venus-IN variant was excluded.

**Figure 3:**
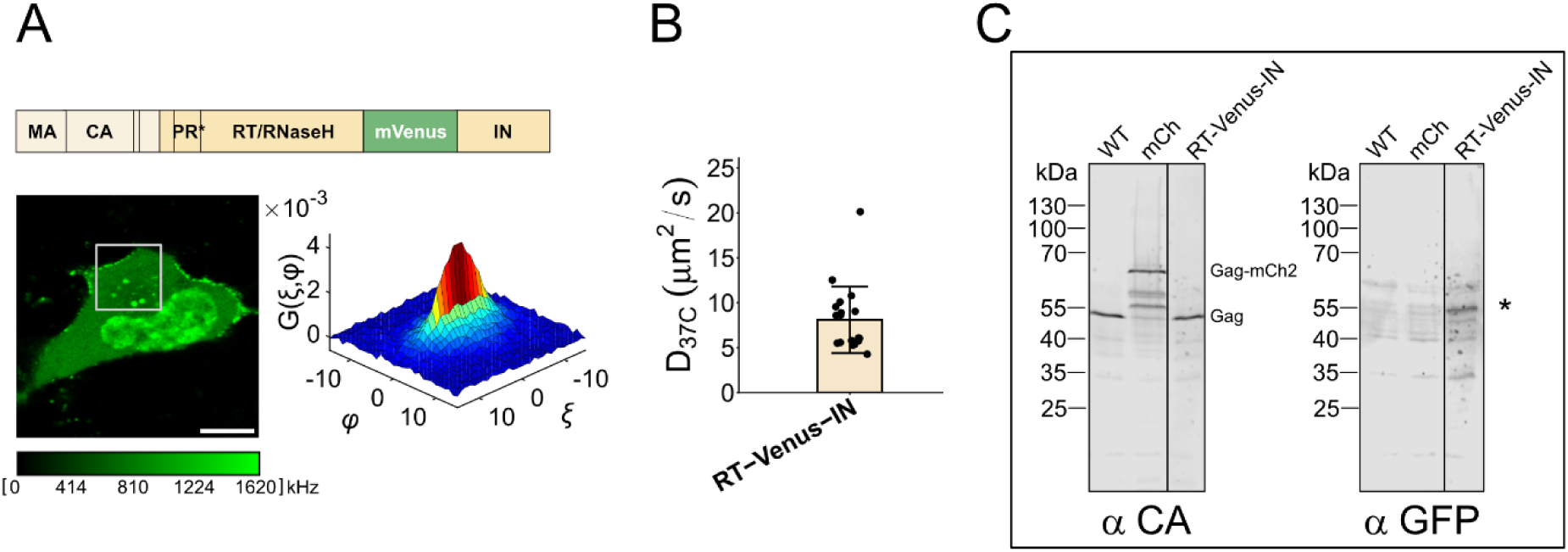
Labeling of GagPol around the IN domain Labeling of GagPol around the IN domain. **A)** Left: Representative image of HeLa cells transfected with the RT-Venus-IN construct. RICS was performed by imaging the ROI highlighted by the white square, and the SACF is reported on the right. Scale bar 10 µm. The schematic of the construct is shown above. **B)** Diffusion coefficients (mean ± standard deviation) of GagPol constructs RT-Venus-IN shown in panel A. Results from 19 individual cells, pooled from two biological replicates, are shown as individual data points. **C)** Immunoblot analysis of GagPol derivative RT-Venus_IN in cell lysates. Cells were transfected with a plasmid expressing wild-type or tagged GagPol in the context of pCHIV (20). At 48h post transfection cell lysates were harvested, proteins were separated by SDS-PAGE and transferred to a nitrocellulose membrane. Gag, GagPol and derivatives were detected by Licor immunoblot using the indicated antisera. Positions of the molecular mass marker are indicated (in kDa). The * indicates the aberrant GagPol product. The immunoblot was spliced to remove the middle lane, and the splice is indicated by a black line.

We next tested a previously generated, unpublished GagPol variant (DL-IN-mCherry), with mCherry fused at the C-terminal domain of IN, and mVenus inserted at the MA/CA junction of Gag. Although C-terminal labeling is a widely utilized strategy, (32–35), prior studies showed that C-terminal tagging of IN might impair GagPol functionality (28). Nevertheless, we tested it as part of this systematic evaluation of labeling strategies. HeLa cells expressing DL-IN-mCherry were imaged 18h post-transfection and displayed a cytoplasmatic fluorescence signal for Gag.mVenus, as expected (27, 30, 31) (Supplementary Figure S3A). In contrast, a homogeneous signal in both cytosol and nucleus was detected for GagPol.mCherry suggesting proteolytic cleavage of mCherry and/or the production of an aberrant, non-full length GagPol. This hypothesis is further supported by the diffusion coefficients extracted by RICS analyses. Specifically, GagPol.mCherry exhibited a diffusion coefficient of 67 ± 9 μm^2^/s (Supplementary Figure S3B, Supplementary Table S2). Although this value may be overestimated due to photobleaching, it is too high to be consistent with full length GagPol, indicating that this construct not usable to image GagPol. In contrast, Gag.mVenus showed a *D* of 4.1 ± 1.3 μm^2^/s, in line with reported values (27). Taken together, these results suggest that placing a fluorescent protein in proximity to the IN domain destabilizes GagPol and leads to partial cleavage by cellular proteases. Therefore, this position was not deemed suitable and was not further pursued.

### Labeling of GagPol by replacing the IN domain with mVenus

Given that GagPol interdomain positioning of mVenus caused aberrant expression and instability of the tagged polyproteins as well as C-terminal labeling, we decided to replace the IN domain with mVenus (ΔIN-Venus). Although the IN domain has functions besides its enzymatic activity in promoting efficient reverse transcription and in morphogenesis of the mature virion (reviewed in (36)) it is not expected to have a significant influence during the early stages of assembly (37). Due to the similar molecular weight of mVenus and IN (∼27 and ∼32 kDa, respectively) the overall polyprotein size remains similar. This also allowed us to insert mCherry2 as a second fluorophore between MA and CA to generate a double-labeled construct (DL-ΔIN). mCherry2 is known for its high brightness, fast maturation rate and high maturation efficiency (34).

Cells co-expressing ΔIN-Venus and pCHIV-mCherry2, as well as cells transfected with DL-ΔIN, displayed a cytosolic GagPol signal colocalizing with Gag (Figure 4A-B). In both cases, the fluorescence signal of GagPol.mVenus was substantially dimmer compared to the previously tested (and discarded) variants. This phenotype is consistent with the expected expression level of GagPol, which is produced by a ribosomal frameshift occurring only in approximately 5% of the translation events (7). This observation supported our assumption that the previously tested tagged variants were not representative of full-length GagPol but instead reflected aberrant, truncated products.

**Figure 4).**
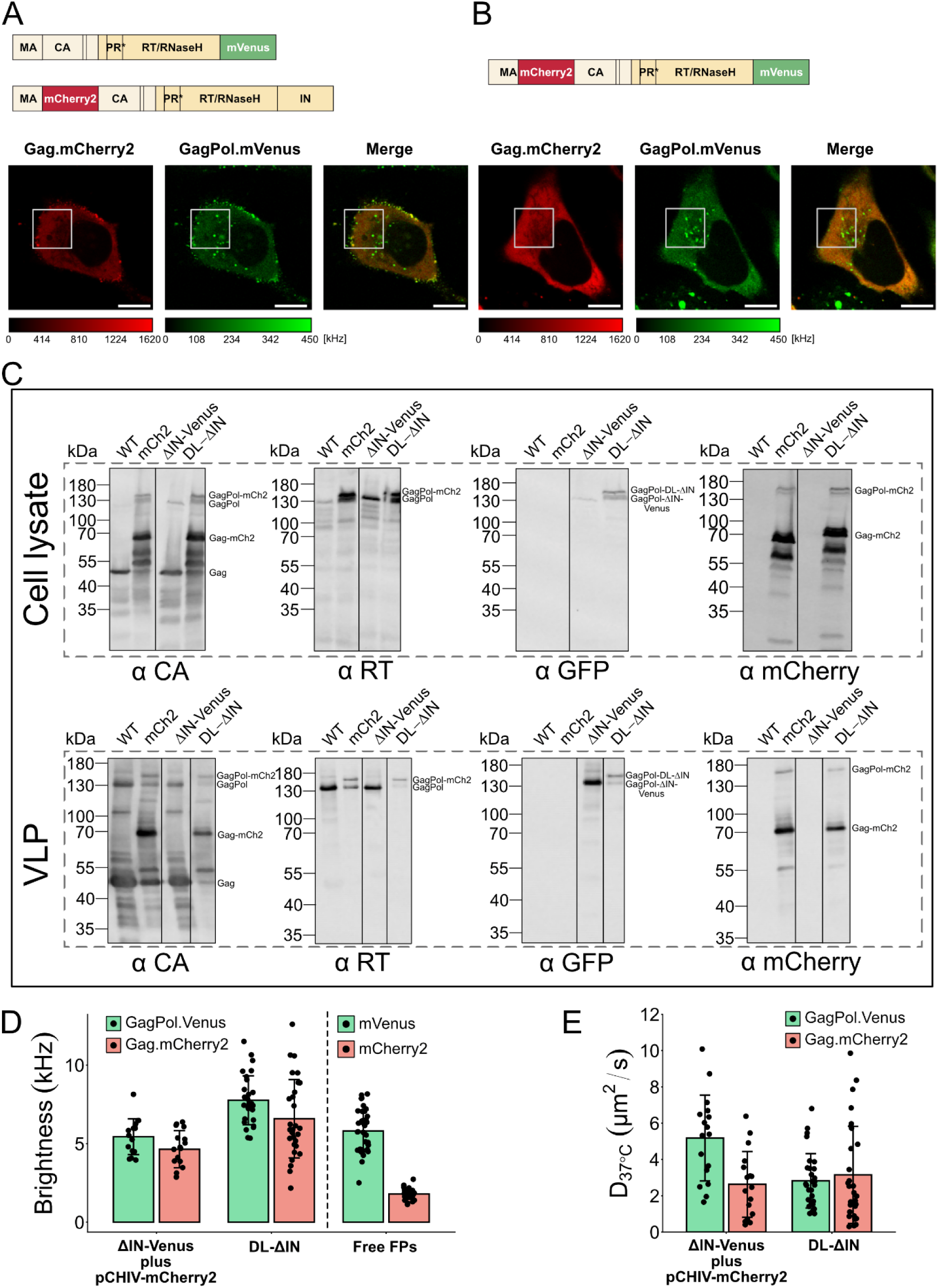
Results for GagPol ΔIN constructs. **A,B**) Representative images of HeLa cells transfected with (A) ΔIN-Venus plus pCHIV-mCherry2 or (B) DL-ΔIN constructs. RICS was performed by imaging the ROI highlighted by the white square. Scale bar 10 µm. **C**) Immunoblot analysis of GagPol derivatives. Cells were transfected with a plasmid expressing wild-type or tagged GagPol together with the other HIV proteins (except Nef) and harvested 48h post transfection. The supernatant was collected for VLPs purification, while the cells were lysed. Proteins were separated by an SDS-PAGE and Gag, GagPol and derivatives were detected by ECL with antisera against CA, RT, Venus and mCherry. mCh, corresponding to pCHIV-mCherry2, is complemented with wild-type at a 1:1 ratio for particle production. Individual immunoblots are outlined by a black border. **D**) Molecular brightness (mean ± standard deviation) of Gag.mCherry2 and GagPol.mVenus obtained from the RICS analyses of the ΔIN-Venus plus pCHIV-mCherry2 (panel A) and DL-ΔIN (panel B) constructs. As a control, the brightnesses of free mVenus and free mCherry2 are reported to the right of the graph. Results from individual cells (n ≥ 17), pooled from at least three biological replicates, are shown as individual data points. **E**) Diffusion coefficient (mean ± standard deviation) of Gag.mCherry2 and GagPol.mVenus obtained from the ΔIN-Venus plus pCHIV-mCherry2 (panel A) and DL-ΔIN (panel B) constructs. Results from individual cells (n ≥ 17), pooled from at least three biological replicates, are shown as individual data points (see Supplementary Tables S1 and S2 for more details).

ΔIN-Venus expressing cells analyzed by immunoblot (Figure 4C) showed a similar profile to wild-type HIV expressing cells, confirming that replacing IN with the fluorescent protein did not significantly impact the integrity of GagPol. The labeled GagPol band of ∼160 kDa was also detected using an anti-GFP antibody, validating fusion with mVenus. Similarly, immunoblot analyses of cells expressing DL-ΔIN revealed a double-labeled GagPol band at ∼180 kDa (187kDa calculated mass), displaying a profile similar to that of pCHIV.mCherry2 (Figure 4C). Therefore, VLP assembly and release was apparently not affected by the presence of both labels as Gag/GagPol ratios look similar to that of pCHIV.mCherry2.

These results validated both ΔIN-Venus and DL-ΔIN as suitable candidates to study the cytosolic behavior of GagPol with fluorescence correlation methods. Due to the low expression levels of GagPol (7), the fluorescence signal in the mVenus channel was close to background levels. For this reason, the single-labeled GagPol was not visualized alone, but cells were always co-transfected with pCHIV-mCherry2. The strong fluorescence signal of Gag.mCherry2 served as a reference to select transfected cells and allowed to probe intermolecular interaction of mVenus labeled GagPol with Gag and GagPol labeled with mCherry2. To validate that the green signal (Figure 4A-B) is specific to labeled GagPol, we performed a RICS analysis on cytosolic ROIs. With this method, we can not only determine the diffusion coefficient of the observed molecule, but also its apparent molecular brightness (*ε*). The RICS analyses showed an average brightness of *ε* = 5.4 ± 1.1 kHz for ΔIN-Venus and of *ε* = 7.8 ± 1.6 kHz for DL-ΔIN, both comparable to the brightness of free mVenus (5.8 ± 1.3 kHz) (Figure 4D and Supplementary Table S1), a value significantly higher than that of autofluorescence (0.6 ± 0.7 kHz, Supplementary Figure S4 and Supplementary Table S1). It should be noted that the cytoplasmatic spots observed in the green channel (Figure 4A-B) were due to autofluorescence and were excluded from the RICS analysis by intensity thresholding. Due to the low expression levels of GagPol.mVenus, these spots become visible and cells transfected with unlabeled pCHIV showed a similar phenotype (Supplementary Figure S4). The interpretation of these signals as autofluorescence is supported by the distinct, shorter fluorescence lifetime profile of these features compared to the one of free mVenus. Cells expressing either the ΔIN-Venus or the DL-ΔIN construct show a long lifetime component in the cytosol, i.e. GagPol.mVenus, and a short lifetime in correspondence to the bright aggregates, i.e. autofluorescence (Supplementary Figure S5).

By analyzing Gag.mCherry2 for both pCHIV-mCherry2 and DL-ΔIN, we observed an average brightness around 6 kHz whereas the average brightness of free mCherry2 was 1.8 ± 0.3 kHz (Figure 4D). We attribute this to the self-assembling property of Gag into slow-moving oligomers in the cytosol, as reported for Gag.mVenus (27). Interestingly, GagPol had a diffusion coefficient of 5.2 ± 2.4 and 2.8 ± 1.5 µm^2^/s for the single and double labeled variants, respectively, which was similar or faster than Gag. In line with previously reported data (27), the diffusion coefficient of Gag was 3.1 ± 2.7 and 2.1 ± 1.5 µm^2^/s for Gag.mCherry2 expressed from DL-ΔIN and pCHIV.mCherry2, respectively (Figure 4E and Supplementary Tables S1 and S2).

### Quantification of the Gag:GagPol ratio via cross-correlation

The established system allowed us to investigate the ratio and potential intermolecular interactions of Gag and GagPol on a single-cell level by cross-correlation RICS (ccRICS). For this, HeLa cells were transfected with either DL-ΔIN or with both ΔIN-Venus and pCHIV-mCherry2. We selected a cytoplasmatic ROI and acquired a series of images in two channels (for mVenus and mCherry2) simultaneously using pulsed interleaved excitation (PIE) (38, 39) in order to minimize the spectral crosstalk that would lead to false-positive cross correlations. Similar to one color RICS, the correlation is then calculated per frame and then averaged over the movie (Figure 5A) to generate the spatial cross-correlation function (SCCF). However, in this case, each image in one channel (mVenus) is correlated with the corresponding frame of the second channel (mCherry2). In contrast to the SACF, the amplitude of the SCCF is proportional to the concentration of cross-correlating molecules, i.e. of double-labeled complexes (38, 39).

**Figure 5:**
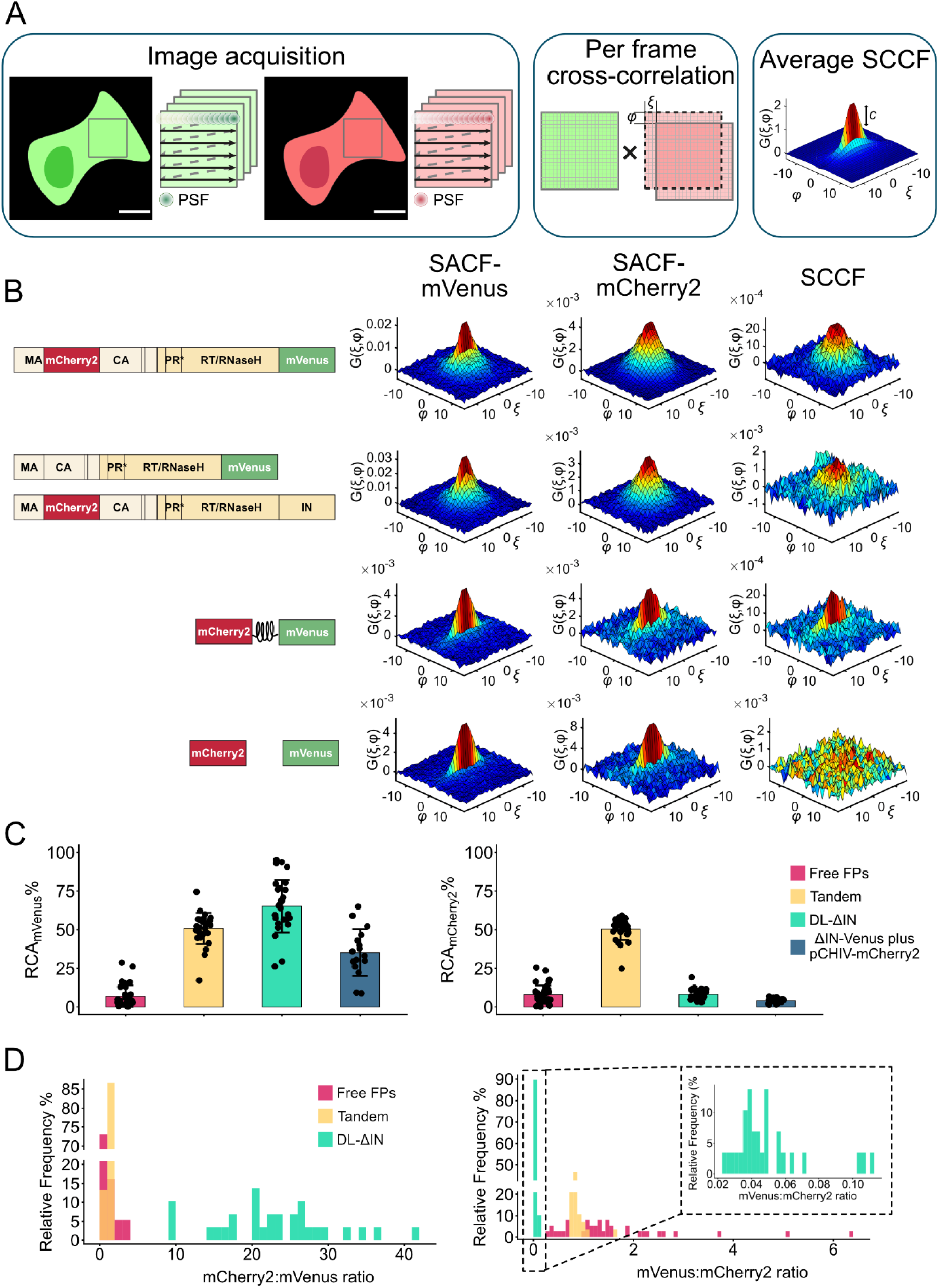
CcRICS analysis of Gag:GagPol interactions and expression ratio. **A)** Left panel: Schematic representation of the cross-correlation RICS workflow. A 12 × 12 µm region of interest is selected and 150 frames are acquired simultaneously in two detection channels. The scanning point spread function (PSF) of the laser is indicated. Central panel: Each frame of the first channel is correlated with the corresponding frame of the second channel, shifted by a spatial or pixel lag (ξ, ψ). This process is repeated for every combination of (ξ, ψ) Right panel: the spatial correlation functions obtained from individual frames were averaged together to obtain the average spatial cross-correlation function (SCCF). **B)** Representative SACFs (SACF.mVenus, SACF.mCherry2) and SCCFs obtained for the variants depicted on the left side: DL-ΔIN, ΔIN-Venus plus pCHIV-mCherry2, mCherry2-mVenus tandem (positive control) and free mCherry2 and mVenus (negative control) from top to bottom. **C)** Relative Cross-correlation Amplitudes (RCA) with respect to the total number of mVenus-labeled molecules (RCA_mVenus_, left) and mCherry2-labeled molecules (RCA_mCherry2_, right) for the indicated variants. Results from individual cells (n ≥ 17), pooled from at least three biological replicates, are shown as individual data points superimposed on bar graphs representing the mean ± SD. **D)** Frequency distribution of the N_corrected_ ratio of mCherry2-over mVenus- labeled proteins (left panel) and vice versa (right panel) for the indicated variants. (see Supplementary Tables S1 and S2 for more details).

We first imaged cells co-expressing free mVenus and free mCherry2 as a negative control and cells expressing a tandem mCherry2-mVenus fusion construct as a positive control. As expected, only a negligible cross-correlation amplitude was detected for the negative control (Figure 5B, first row) whereas a significant positive SCCF amplitude was detected for the tandem construct (Figure 5B, second row). This confirms that mCherry2 and mVenus are co-diffusing and provides a measure of the expected maximum cross-correlation amplitude. The ccRICS analyses performed on cells transfected with the DL-ΔIN construct showed a clear positive cross-correlation (Figure 5B, third row) verifying that GagPol was indeed double-labeled. Interestingly, also cells co-expressing ΔIN-Venus and pCHIV-mCherry2 exhibited positive cross-correlation amplitudes (Figure 5B, last rows), suggesting inter-molecular associations of mVenus-labeled GagPol with Gag or other GagPol molecules labeled with mCherry2 and produced by pCHIV-mCherry2.

To determine the cytosolic Gag:GagPol ratio, we calculated the relative cross-correlation amplitudes (RCA), i.e. the ratio of the SCCF amplitude to the SACF amplitude of mVenus (RCA_mVenus_) or mCherry2 (RCA_mCherry2_) (Figure 5C). Under simplifying assumptions (see Material and Methods for details), RCA_mVenus_ estimates how many double-labeled complexes exist compared to the total number of mVenus-labeled molecules, and RCA_mCherry2_ calculates the fraction of double-labeled complexes with respect to mCherry2 containing proteins. Using free fluorescent proteins as a negative control, we established a residual baseline of 7 and 8 % for RCA_mVenus_ and RCA_mCherry2_ respectively (Supplementary table S3). Although PIE can remove spectral crosstalk, spatial correlations within cells, the presence of autofluorescence and spurious fits to the noisy correlation data can still lead to a non-zero cross-correlation amplitude. For the tandem fusion protein with mCherry2 and mVenus connected by a 19 amino acids linker, we have a RCA of 50 %, a typical value for fluorescent proteins (40–42). Theoretically, for a fusion protein with a 1:1 stoichiometry, one would expect an RCA of 100%. However, multiple factors can reduce the amplitude including different maturation yields of the fluorescent proteins, imperfect overlap of different observation volumes, different observation volume sizes, Förster Resonance Energy Transfer (FRET) and photobleaching (40). The similarity between the RCA_mVenus_ and RCA_mCherry2_ values suggests that the maturation rates of mVenus and mCherry2 are similar but does not exclude that a fraction of the fluorophores might be in a dark state and/or FRET might be occurring between the two fluorescent proteins, despite the linker between them.

The DL-ΔIN construct showed a RCA_mVenus_ of 65 ± 17 %, in essence confirming the presence of both mCherry2 and mVenus within GagPol. The fact that this value is higher than our positive control can most likely be attributed to the formation of oligomers and hence a different stoichiometry ratio than 1:1 (43). This observation is supported by the increased brightness of Gag.mCherry2 compared to the monomeric fluorescent protein (Figure 4D), which suggests the formation of oligomers containing more than one Gag molecule. Conversely, since the brightness of GagPol.mVenus is comparable to that of monomeric mVenus (Figure 4D), only one GagPol molecule may be part of the complex. Interestingly, the analysis of cells co-expressing ΔIN-Venus and pCHIV-mCherry2, in which only GagPol inter-molecular interactions are probed (with either Gag or other GagPol), revealed a RCA_mVenus_ of 35 ± 15 %. When considering the maximum cross-correlation amplitude obtained for the DL-ΔIN construct (65 %), this would suggest a significant amount of GagPol being associated with Gag through inter-molecular interactions. The RCA_mCherry2_ values, 4.1 ± 1.7 % for the co-expressed single labeled plasmids and 8 ± 6 % for DL-ΔIN (Figure 5C, right panel, and Supplementary table S3), further corroborate the hypothesis that the Gag-GagPol interactions are divided between inter- and intra-molecular interactions. It should be noted that a low RCA_mCherry2_ is expected because of the lower expression levels of GagPol (7, 9, 44). However, as the residual cross-correlation amplitudes in our control measurements were approximately 7%, we are close to the detection limit and interpretation should be made with caution. Taken together, these results suggest that approximately half of the expressed GagPol molecules exist in a monomeric form, while the remaining half participates in the formation of small oligomers containing Gag, which have been previously hypothesized to be cytosolic nucleation site for assembly (27).

To evaluate the relative abundance of Gag and GagPol in individual cells, we use the ratio of the average number ( *N*) of the respective proteins. Due to various challenges of a quantitative correlation analysis, we calculate *N* from the intensity normalized to the molecular brightness of a monomeric fluorescent label (see Materials and Methods). By analyzing the frequency distribution of these ratios (Figure 5D), we first confirmed that the mCherry2-mVenus tandem protein has the expected 1:1 stoichiometry. The ratio of the co-expressed free proteins was still centered around 1:1 but spread more widely due to expression from co-transfected plasmids. Cells transfected with the double-labeled DL-ΔIN variant revealed an average Gag:GagPol ratio of 23 ± 8 (Supplementary table S3), consistent with the GagPol expression ratio derived from bulk analyses (7, 9, 10). However, the wide spread of the distribution (Figure 5D, left panel) indicates considerable heterogeneity at the single-cell level. Likewise, the ratio of GagPol:Gag was determined to be 0.050 ± 0.022, on average (Supplementary table S3), with a very disperse distribution (Figure 5D, right panel). We also calculated the average Gag:GagPol ratio for ΔIN-Venus co-expressed with pCHIV.mCherry2, revealing an average value of 26 ± 12 (Supplementary table S3).

Overall, these results indicate that, while the average Gag:GagPol ratio is consistent with ensemble biochemical studies, individual cells show different relative expression rates. This highlights the importance of single-cell studies, which are essential to capture cellular heterogeneity. Knowing that an improper Gag:GagPol ratio is detrimental for HIV-1 fitness (10), identifying the cellular factors that influence the expression rate is crucial for understanding the infectivity mechanisms.

## Discussion

Despite the critical role of viral-enzyme incorporation for HIV-1 infectivity, GagPol oligomerization during its trafficking towards the plasma membrane (i.e. in the cytosol) and its dynamics during viral assembly are not fully understood. To address this at the single-cell level, we systematically tested different approaches for fluorescently labeling GagPol in the context of a full-length HIV-1 construct comprising all viral proteins except Nef, thereby maintaining a native context for assembly. To evaluate virion assembly independently of maturation, all the investigated variants carried a catalytically inactive protease. First, we labeled GagPol by inserting a fluorescent protein at the PR-RT junction, a strategy previously adopted by Takagi et al. (24). The authors used a frameshift mutant of GagPol to express GagPol only and assayed GagPol homodimerization with a FRET assay. However, the stability of labeled GagPol was not assessed in their work. While the fluorescent phenotype obtained by our PR-Venus-RT variant was consistent with Takagi et al. (Figure 2B), detailed immunoblot analyses revealed the production of a dominant, aberrant polypeptide containing RT and mVenus but not CA (Figure 2D, Supplementary Figure S2). Despite exhibiting a Gag expression profile similar to that of wildtype, the expression of this aberrant product impaired the use of this variant to monitor GagPol. The interaction previously observed for this construct via FRET could be potentially explained even for the aberrant product by dimerization of RT (45).

Labeling GagPol at its C-Terminus or at the N-terminus of IN was also not suitable for assembly studies, as it led to unspecific cleavage of the fluorescent tag (Figure 3, Supplementary Figure S3). GagPol labeled at the N-terminus of IN was previously reported by Mamede et al. to be incorporated into pseudo-typed VSVG particles that partially maintained viral infectivity. This suggested that, once packaged, GagPol remained functional to some extent (28). While this approach could be a useful tool to fluorescently tag VLPs, the cytoplasmic stability of GagPol was not investigated in their study. Our results indicate unspecific cleavage of mVenus by cellular proteases, but the findings of Mamede et al., may suggest at least part of the GagPol expressed by the RT-Venus-IN variant might retain its functionality. Nevertheless, the presence of cleaved mVenus-labeled constructs hampers the use of this construct to investigate the cytosolic dynamics of full length GagPol.

Therefore, since inter-domain and C-terminal tagging of GagPol altered the expression and/or stability of the polyprotein, we finally labeled GagPol by replacing the IN domain with mVenus (ΔIN-Venus). IN has been shown to affect mature morphogenesis but is not expected to have a significant influence during the early stages of assembly (37, 46, 47). This strategy resulted in the expression of a stable GagPol variant that was incorporated into viral particles and displayed a cytosolic fluorescent signal (Figure 4) that can be used to follow GagPol assembly dynamics. Furthermore, the stability of full length GagPol was retained even after the addition of a second label, mCherry2, between MA and CA of Gag. Hence, we could also generate a double-labeled variant (DL-ΔIN) where both Gag only and GagPol are visible (Figure 4). We used the DL-ΔIN variant, in which Gag and GagPol are produced according to their wildtype frameshift efficiency, to quantify the relative abundance of Gag and GagPol in single cells. We used the molecular brightness of free mVenus and mCherry2 to estimate the relative abundance of Gag and GagPol. The results showed a Gag:GagPol ratio on average close to 23:1, in large agreement with previous studies (7, 9, 10) but also revealed a high cell-to-cell variability of Gag and GagPol stoichiometry (Figure 5D).

RICS analyses performed in cells expressing the mVenus-labeled GagPol (from either ΔIN-Venus or DL-ΔIN) and mCherry2-labeled Gag showed that GagPol exhibits a lower tendency to oligomerize compared to Gag at their respective concentrations. This observation was supported by the molecular brightness and diffusion coefficient of the polyproteins. The average brightness of GagPol was comparable to that of monomeric mVenus, while the average brightness of Gag.mCherry2 was approximatively threefold higher than monomeric mCherry2 (Figure 4D). This suggests that GagPol does not interact with other GagPol molecules for the conditions measured, while Gag tends to form small multimers, in line with previously reported data (27).

The diffusion coefficient of GagPol (2.8-5.2 µm^2^/s) was comparable to that of Gag (2.1-3.1 µm^2^/s) (Figure 4E), despite the large difference in molecular weight (160 and 187 kDa for the single and double labeled GagPol variants, respectively, and 82 kDa for Gag.mCherry2). If Gag and GagPol were present as individual proteins without interactions, we would expect Gag to diffuse faster than the larger GagPol. However, the interaction of Gag with RNA and cellular proteins has been shown to reduce the mobility of Gag compared to a freely diffusing protein of equivalent size (27, 48). Thus, the comparable diffusion coefficients of Gag and GagPol suggest that factors beyond molecular weight influence their mobility and may suggest that GagPol is less prone to interact with RNA (49) or cellular components. However, since diffusion depends on hydrodynamic radius rather than molecular weight alone, conformational differences could also influence their mobility but were not investigated in this study.

The cross-correlation analysis of GagPol and Gag showed that GagPol is involved in inter-molecular interactions with Gag. Indeed, a positive cross-correlation was detected in cells co-expressing ΔIN-Venus and pCHIV-mCherry2 (Figure 5B-C). Although pCHIV-mCherry2 produces both Gag and GagPol labeled at the MA/CA junction, it is reasonable to speculate that the detected cross-correlation primarily reflects interactions between GagPol.mVenus and Gag.mCherry2 rather than between GagPol molecules. This interpretation is supported by the fact that GagPol dimerization would lead to a twofold increase in molecular brightness compared to monomeric mVenus, which was not observed for either ΔIN-Venus nor DL-ΔIN (Figure 4D). Similarly, a positive cross-correlation was detected in cells expressing double-labeled GagPol (DL-ΔIN), where the relative cross-correlation amplitude exceeded that of the positive control, a mCherry2-mVenus tandem protein (Figure 5B-C). This typically occurs when the observed interaction has a different stoichiometry ratio than 1:1 (43), and is therefore consistent with the formation of small Gag-GagPol oligomers. Taken together, these results suggest that GagPol is involved in intermolecular association with Gag, forming small oligomers composed of one GagPol and several Gag molecules.

In summary, in this study, we showed that, by replacing IN with a fluorescent protein, we were able to obtain a stable GagPol variant suitable for studies of early HIV assembly, while inserting a fluorescent tag within the Pol region or at the C-terminus caused protein instability and aberrant products. The systematic investigation of various labeling positions highlighted that controls from multiple perspectives were necessary. Simple western blots and fluorescence phenotypes are not always sufficient. Rather, detailed immunoblots together with direct monitoring of the label is required. Overall, these labeled variants pave the way for detailed investigations of GagPol during HIV assembly to shed light on cytoplasmic trafficking and assembly kinetics.

## Materials and Methods

### Plasmids

All HIV-1 plasmids utilized in this study are based on pCHIV, a non-replication competent proviral derivative of HIV-1 NL4-3, lacking the long terminal repeat regions and the *nef* gene (20). Plasmid pCHIV-mCherry2 was generated by replacing the mCherry sequence of pCHIV-mCherry, inserted between MA and CA region of *gag* ORF (20) with a PCR fragment encoding mCherry2 expanded from Addgene plasmid #54563, using NEBuilder HiFi DNA Assembly (#E2621S). The RT-Venus-IN was generated by inserting a PCR product encoding for mVenus, amplified from pmVenus-C1 plasmid (27), in a pCHIV derivative with a unique KpnI site at the RT-IN junction within the *pol* ORF. Similarly, a pCHIV derivative with unique KpnI site at the PR-RT junction within the *pol* ORF was used to generate the plasmids PR-Venus-RT, PR PR-FL-Venus-RT, PR-RL-Venus-RT and PR-CFP-RT. The inserts were expanded by PCR and inserted via KpnII digestion. Each respective insert was amplified by PCR, with the linker (FL, RL) included in the sequence of the PCR primers. The eCFP sequence was obtained from the peCFP plasmid purchased from Invitrogen (Carlsbad, CA, USA).

Plasmid ΔIN-Venus was obtained by replacing the IN sequence with a PCR fragment encoding mVenus (from pmVenus-C1 plasmid) using NEBuilder HiFi DNA Assembly (#E2621S). The double labeled plasmid DL-ΔIN was generated by replacing a fragment of ΔIN-Venus flanked by ApaI sites (comprising the 5’ Gag till the NC domain) with the corresponding fragment of pCHIV-mCherry2. All HIV-1 derived plasmids were inactivated for protease activity by a D25N amino acid substitution in the PR domain, achieved through site directed mutagenesis (primers: fwd gcaattaaaggaagctctattaaacacaggagcagatg, rev atcatctgctcctgtgtttaatagagcttcctttaattgc).

Free monomeric mVenus was expressed from plasmid pmVenus-C1 (39) and a plasmid encoding free monomeric mCherry2 was purchased from Addgene (#54563). The plasmid encoding the mCherry2-mVenus tandem was generated by inserting mVenus sequence amplified by PCR into the mCherry2 plasmid via NEBuilder HiFi DNA Assembly (#E2621S). All the primers used for molecular cloning were purchased from Eurofin and are listed in Supplementary Table S4. The sequences of the plasmids used in this study was verified by whole plasmid sequencing and are available on request.

### Cell culture and transfection

HeLa and HEK293T cells were cultivated in high glucose Dulbecco’s Modified Eagle Medium (DMEM) supplemented with Glutamax™ (Gibco™, #61965026), 10% fetal calf serum (Gibco™, #26140079, USA origin) heat-inactivated for 30 min at 56 °C and 1 % Penicillin-Streptomycin (Gibco™, #15140-122), at 5% CO_2_ and 37°C in a humidified atmosphere.

For microscopy experiments, 5 × 10^4^ cells per well were seeded in complete medium in 8-well coverslips (Nunc™ Lab-Tek™ II Chambered Coverglass, Thermo Fisher Scientific). 24h post-seeding, 2 µg of plasmid DNA were mixed with 6 µl of FuGENE® 4K Transfection Reagent (Promega, #E5911) and Opti-MEM™medium (Gibco™, # 31985062) in a final volume of 100 µl and incubated for 15 minutes at room temperature. For the HIV-1 plasmid 200 ng plasmid were added to each well. For plasmids expressing fluorescent proteins, 50 ng were transfected. For co-transfection experiments, 1:1 molar ratio was used.

For viral particle production, ∼5×10^6^ HEK293T cells were seeded in T175 flasks (Sarstedt, #89.3912.002). 24h after seeding, the HEK293T cells were transfected with 70 µg of plasmid DNA, diluted in 3.5mL sterile 250mM CaCl_2_, followed by the addition of 3.5mL sterile 2X HBS buffer (280mM NaCl, 50mM HEPES, 1.5mM Na_2_HPO_4_ hydrate, pH 7.1). The mixture was vortexed and incubated for 20 minutes at room temperature before adding to the cells medium. 4 hours later, the medium was replaced. For cell lysates experiments, 2.6×10^5^ HEK293T cells were seeded in T25 flasks (Sarstedt, #83.3910.002) and transfected similarly with 5µg of plasmid DNA, followed by the addition of 1mL of 2X HBS buffer and 1mL of CaCl2.

### Fluorescence imaging

At 15–18 h post transfection, the cell culture medium was replaced with phenol red free DMEM FluoroBrite™ (Gibco™, Thermo Fisher Scientific). Imaging was performed at 37 °C on a home-built confocal laser scanning microscope (CLSM) (Supplementary Figure S6) as described elsewhere (39). A 100× oil immersion objective (Apo-TIRF 100x Oil/NA 1.49, Nikon) was used for all measurements.

With the exception of the experiments involving the PR-CFP-RT, pulsed lasers of wavelengths 470 nm and 560 nm (LDH-P-C-470 and LDH-P-FA-560, PicoQuant) were used for the excitation of mVenus and mCherry2, respectively. Unless otherwise stated, the 470 nm laser was set at power of 5 μW before the objective, while the 560 nm laser was set at 2 μW, to compensate for the 20-fold higher expression of Gag.mCherry2 compared to GagPol.mVenus, expected due to the Gag: GagPol translation ratio. For consistency, these laser intensities were used for all experiments involving mCherry2 and mVenus. The lasers were synchronized to a pulse repetition of 25 MHz and delayed of ∼17 ns with respect to each other, to achieve pulsed interleaved excitation (PIE) (38). Using PIE allows minimization of crosstalk and hence remove false positive cross-correlations. The fluorescence emission was recorded by avalanche photodiode detectors (APDs) (Count® Blue, Laser Components, after a 520/40 nm bandpass filter for mVenus and SPCM-AQR-14, PerkinElmer, after a 595/50 bandpass emission filter for mCherry2). The alignment of the microscope was routinely checked by measuring a 10 nM mixture of ATTO488-COOH and ATTO565-COOH in dPBS (Gibco™). Given the known diffusion coefficients of these fluorophores, a Fluorescence Correlation Spectroscopy (FCS) analysis is used to determine the size of the observation volume (50).

To image the PR-CFP-RT construct, a single 440 nm pulsed diode laser (LDH-P-C-440, PicoQuant) was used at a laser power of 2 µW before objective and a repetition rate of 40 MHz. Fluorescence emission was collected by the Count® Blue detector after a 480/40 nm emission filter. The alignment was evaluated by FCS measurement of ATTO425-COOH in dPBS.

For all experiments, the excitation and emission were synchronized via time-correlated single-photon-counting electronics (TCSPC cards, Becker SPC-150 and Hickl). For every cell, a 60 × 60 µm image was recorded as a reference, followed by the acquisition of a 12 × 12 µm region of interest (ROI) positioned in the cell cytoplasm. A total of 150 frames were acquired, with a frame time *τ_f_* = 1 s, line time *τ_l_* = = 3.33 ms, pixel dwell time *τ_p_* = 11.11 μs, and pixel size *δ_r_* = 40 nm. The raw photon data stream acquired by the APDs was converted into images with the software PIE Analysis in MATLAB (PAM) (51). To simplify the localization of successfully transfected cells, a widefield unit is integrated into the CLSM. Switching from widefield to confocal (and vice versa) was possible by exchanging a mirror present in the cube slider of the microscope body. This did not affect the alignment of the confocal path.

### Raster Image (Cross)Correlation Spectroscopy (RICS and ccRICS)

RICS and ccRICS analyses on the 12×12 µm image series were performed with the Microtime Image Analysis (MIA) submodule of PAM (51). Slow fluctuations and cellular movements were removed pixel-wise by applying a three-frame moving average correction as previously described (27). Subsequently, a frame-wise intensity thresholding was applied to exclude spatial inhomogeneities, such as bright aggregates, but also to exclude background surrounding the cell, in case the 12 × 12 µm ROI does not completely lie within the cytoplasm. The spatial auto- and cross-correlations are then calculated for each individual frame using the Arbitrary RICS algorithm previously developed (52), which allows to correlate non-squared ROIs.

The average SACFs and SCCFs were analyzed with a two-component model, accounting for a mobile species and an immobile one, to be consistent with previously published results (27). The used model assumes Brownian diffusion and a 3D-Gaussian focal shape (accounted for with the introduction of the geometrical factor *γ* = 2^−3/2^) and the correlation function *G*(*ξ*, *ψ*) corresponds to:

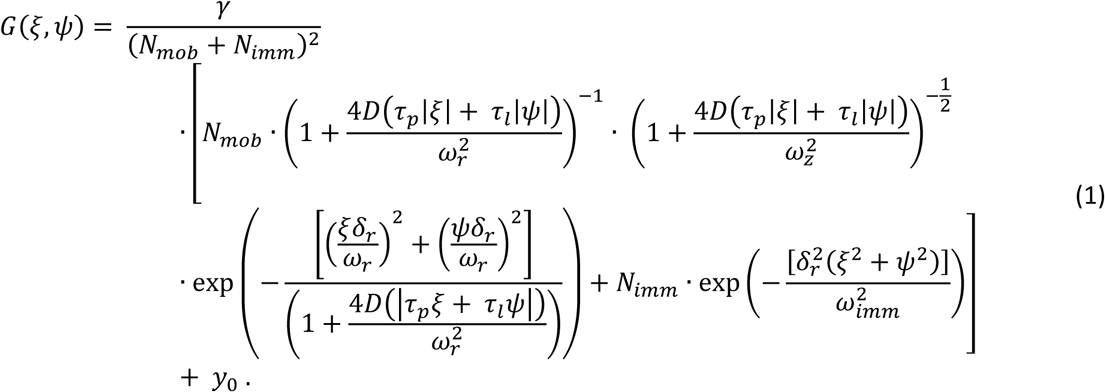

Here, *N_mob_* and *N_imm_* represent the average number of mobile and immobile fluorescent molecules in the observation volume, characterized by a lateral and axial radius equal to *ω_r_* and *ω_z_*, respectively. To account for the different refractive index of the cell cytoplasm compared to the dPBS calibration reference, the axial radius of the observation volume was estimated by globally linking *ω_r_* between SACFs in a large dataset. *ξ* and *ψ* are the spatial correlation lags in the fast and slow scanning axes. The correlation at zero lag time was omitted from analysis due to the contribution of uncorrelated shot noise. *D* is the diffusion coefficient of the mobile species and *ω_imm_* is the half-width of the immobile species 2D Gaussian, at 1/e^2^ of the maximal intensity. *y*_0_ is a baseline correction term. For this work, *N_mob_* and *N_imm_* were summed to a *N_tot_* value specific to each channel, obtaining from the SACFs the total average number of green (*N_G_*) and red (*N_R_*) molecules. The total average number of particles with both green and red fluorescence *N_GR_* was calculated from the apparent number obtained from the SCCF (*N_CCapp_*) according to:

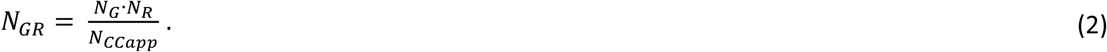

The molecular brightness (*ε*), i.e. the number of photons detected per molecule per second in the excitation volume, is calculated by dividing the average intensity of the acquired image series by *N_tot_*. For cells expressing mVenus-labeled GagPol, cells displaying a molecular brightness lower than 3.8 kHz, corresponding to the mean molecular brightness of free mVenus (5.8 kHz) minus two standard deviations (3.2 kHz), were considered as autofluorescent and discarded from analysis.

The fraction of mobile species (*F_mob_*) is reported in Supplementary Tables S1 and S2 and calculated as:

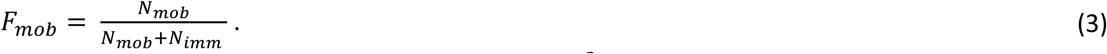

Cells in which one of the SACFs showed a reduced *χ*^2^≥ 3 or with weighted residual mostly above 3, were excluded from the analyses

### Relative Cross-correlation Amplitude (RCA) and *N* ratio calculation

RCA_mVenus_ was obtained by dividing *N_GR_* by *N_G_* by and RCA_mCherry2_ by dividing *N_GR_* by *N_R_* . It should be noted that the quantification of the cross-correlation signal, i.e. the RCA calculation, is performed under the simplifying assumptions of a 1:1 binding stoichiometry and no changes in brightness upon binding (43). While these assumptions hold true for the free proteins and the tandem construct, they are not valid in case of the GagPol constructs, as discussed in the Results section. Therefore, a more quantitative assessment of the GagPol:Gag ratio (and vice versa) was performed by calculating the *N* ratio of mVenus labeled proteins to mCherry2 labeled proteins (and vice versa), after correcting *N_G_* and *N_R_* for potential oligomerization by taking into account the molecular brightness of the monomeric fluorescent proteins. *N_G_*_,*normalized*_ and *N_R_*_,*normalized*_ were obtained by dividing the average intensity of the acquired image series by the molecular brightness of monomeric mVenus and mCherry2, respectively. With this normalization, the ratio of *N_G_*_,*normalized*_ over *N_R_*_,*normalized*_ (and viceversa) was calculated for each individual cell, after which a frequency distribution analysis was performed and shown in figure Figure 5D. Bar graphs and frequency distributions were generated using RStudio: Integrated Development Environment for R. Posit Software, PBC, Boston, MA (Posit team 2025, http://www.posit.co/) and the tidyverse collection of packages (53).

### Absolute concentration calculation

Absolute concentrations (*c*) (Supplementary Tables S1 and S2) were calculated by normalizing the average fluorescence intensity to the molecular brightness of the monomeric reference (mVenus or mCherry2) to cancel the influence of oligomerization on concentration, thereby reporting concentration as monomer equivalents and eliminating contributions from oligomerization. The resulting *N_normalized_* was used to calculate absolute concentrations as follows:

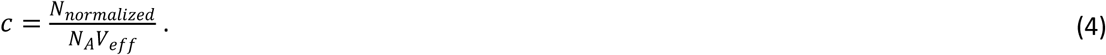

with *N_A_* being the Avogadro’s number, and *V_eff_* being the effective observation volume, corresponding to 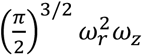.

### Fluorescence lifetime imaging microscopy (FLIM) with Phasor approach

The FLIM-Phasor approach (54) was used to extract the fluorescence lifetime information from the raster scanned images acquired on the CLSM. The raw photon data were analyzed in PAM (51) as previously described (6). Briefly, the pixel-wise lifetime decay is transformed into Fourier space, and the sine and cosine Fourier components of the transformation constitute the phasor coordinates *g* and *s* respectively. An aqueous solution of Atto488-COOH was measured at 37 °C and used as a reference to correct for the instrument response function of the system.

### Virus-Like Particle (VLPs) purification

50 ml supernatant of transfected HEK293T cells was harvested 48h post-transfection, cleared from cellular debris by centrifugation and filtration through a 0.45 µM nitrocellulose membrane and loaded on a 20% (w/v) sucrose cushion in a polyallomer ultracentrifuge tube. The VLPs were concentrated by centrifugation for 1.5 h at 28 000 rpm at 4°C in a Sorvall™ WX ultracentrifuge (Thermo Scientific™) and resuspended in 30µL of PBS.

### Immunoblot

HEK293T were harvested 48h post-transfection: cells were washed with phosphate-buffered Saline (Gibco™^TM^, Fisherscientific), detached with Trypsin, pelleted, resuspended in PBS, pelleted a second timeand resuspended in the lysis buffer (125mM Tri-HCl, pH 6.8, 2% SDS (w/v), 10% glycerin, 5% β-mercaptoethanol, supplemented with protease inhibitor (cOmplete mini™ protease inhibitor cocktail, Roche). Purified VLPs mixed with lysis buffer and lysates were incubated 5’ at 95°C. The denatured proteins were loaded together with the PageRuler Prestained Protein Ladder, 10 to 180kDa (ThermoScientific), as a molecular weight marker on a 12% SDS-polycrylamide gel. SDS-PAGE was performed at a constant current of 50 mA and a maximum voltage of 200V until the dye front reached the bottom of the gel. Three Whatman 3MM paper sheets were soaked in Buffer A2 (25 mM Tris-HCl, 20% methanol, pH 10.4), followed by three sheets soaked in Buffer A1 (300 mM Tris-HCl, 20% methanol, pH 10.4). Next, a nitrocellulose membrane (Amersham Protran, 0.45 µm NC) soaked in Buffer A1 was placed on top of the Whatman stack, followed by the gel soaked in Buffer C (25 mM Tris-HCl, 20% methanol, 40 mM 6-aminocaproic acid, pH 9.4), and finally, three sheets of Whatman paper soaked in Buffer C. The sandwich was assembled in a semi-dry transfer system, and transfer was performed at a constant current of 50 mA for 1 h 30. With the exception of the immunoblots shown in Figure 3C, the membranes were then blocked overnight in blocking buffer (5% w/v milk in PBS with 0.1% v/v Tween 20) at 4°C and then probed for one hour at room temperature with antisera raised against recombinant HIV-1 CA (Sheep, in-house, diluted 1:5000) and RT (Rabbit, in-house, 1:1000) and primary polyclonal antibodies anti-GFP (Abcam, ab290, Rabbit, diluted 1:2000) and anti-mCherry (Abcam, ab167453, Rabbit diluted 1:5000) diluted in blocking buffer. The immunoblots were washed with PBS added with 0.1% Tween20 and completed with a one hour incubation with secondary antibodies Goat Anti-Rabbit HRP (Invitrogen #31460 diluted 1:30 000) and Donkey Anti-Sheep-HRP (Invitrogen #A16041 diluted 1:50 000) diluted in blocking buffer. Membranes were revealed by chemiluminescence (ECL SuperSignal™ Western Blot Substrate Package, Pico PLUS Thermo Scientific™) on a Bio-Rad ChemiDoc™ Imager. For the immunoblots shown in Figure 3C, the membranes were blocked for 20 min at room temperature with Intercept (PBS) Blocking Buffer (LICORBIO) diluted 1:3 in PBS. Subsequently, the membranes were probed overnight at 4°C with antisera raised against recombinant HIV-1 CA (Sheep, in-house, diluted 1:5000) or against recombinant GFP (Rabbit, in-house, 1:1000) diluted in the Intercept Blocking Buffer. After washes with PBS added with 0.1% Tween20, the immunoblots were incubated for 30 min at room temperature with fluorescently labeled secondary antibodies. Detection was performed with an infrared imaging system (LiCOR Odyssey, LiCOR Biosciences) according to the manufacturer’s instructions.

## Acknowledgement

We thankfully acknowledge the financial support of the Deutsche Forschungsgemeinschaft (DFG, German Research Foundation) – Project-ID LA 1971/8-1 and EXC3092/1-533751719 from Germany’s Excellence Strategy, and from the Federal Ministry of Education and Research (BMBF) and the Free State of Bavaria under the Excellence Strategy of the Federal Government and the Länder through the ONE MUNICH Project Munich Multiscale Biofabrication. D.C.L. gratefully acknowledges the financial support of the Ludwig-Maximilians-Universität München via the Department of Chemistry, the Center for NanoScience (CeNS) and the LMUinnovativ program BioImaging Network (BIN). We gratefully acknowledge Prof. Hanna-Mari Baldauf for providing access to the ultracentrifuge for viral particle purification.

