## Supplementary Information for "Developing a fluorescent derivative of GagPol as a tool for live-cell imaging of HIV-1 assembly"

### Supplementary figures

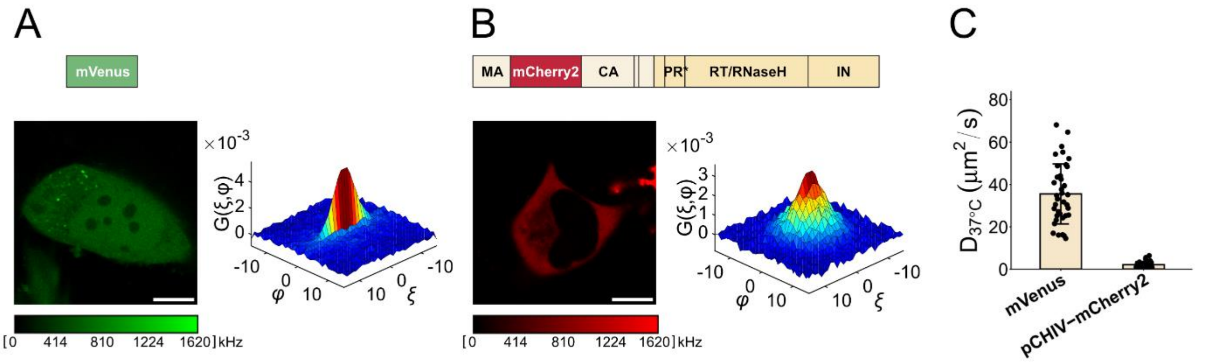

***Figure S1: Control RICS analyses of diffusion. A)*** *Representative image of HeLa cells transfected with* ***mVenus*** *construct (freely diffusing). RICS was performed by imaging a cytoplasmatic ROI of 12 × 12 µm, and the SACF is reported on the right. Scale bar 10 µm.* ***B)*** *Representative image of HeLa cells transfected with* ***pCHIV-mCherry2*** *construct (producing Gag.mCherry2). RICS was performed by imaging a cytoplasmatic ROI of 12 × 12 µm, and the SACF is reported on the right. Scale bar 10 µm.* ***C:*** *Diffusion coefficient (mean ± standard deviation) of free mVenus and Gag.mCherry2 of panel A and B. Results from individual cells (n ≥ 33), pooled from at least three biological replicates, are shown as individual data points (see Supplementary Table S1 and S2 for more details).*

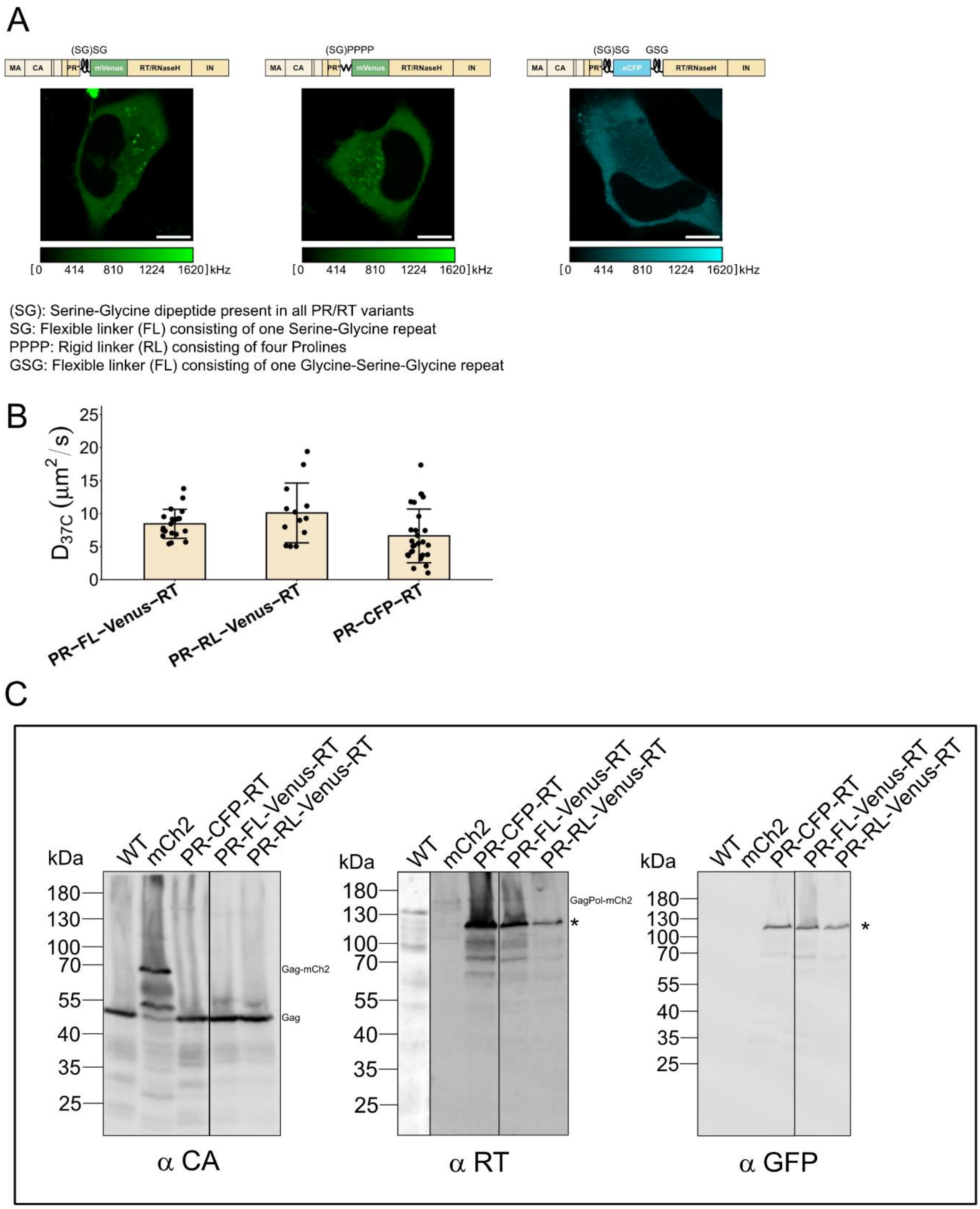

***Figure S2: Effect of different linker and fluorophores on the PR-RT labeling position. A***) *Representative images of HeLa cells expressing the construct depicted above. From left to right:* ***PR-FL-Venus-RT, PR-RL-Venus-RT, PR-CFP-RT****. It should be noted that all PR/RT variants, including the initial PR-Venus-RT, contain a Serine-Glycine dipeptide (SG) upstream of the fluorescent moiety. The PR-CFP-RT variant was imaged with a laser power of 2 µW, measured before the objective.* ***B)*** *Diffusion coefficient (mean ± standard deviation) obtained by RICS analyses of the constructs shown in A. Results from individual cells (n ≥ 13), pooled from at least two biological replicates, are shown as individual data points (see Supplementary Table S1 for more details).* ***C)*** *Immunoblot analysis of GagPol derivatives. Cells were transfected with a plasmid expressing wild-type or tagged GagPol together with the other HIV proteins (except Nef) and harvested 48h post transfection. Cells were lysed, proteins were loaded on an SDS-PAGE and Gag, GagPol and derivatives were revealed by ECL with antisera against CA, RT and Venus. The * indicates the mass of the aberrant GagPol product. Individual immunoblots are outlined by a black border.*

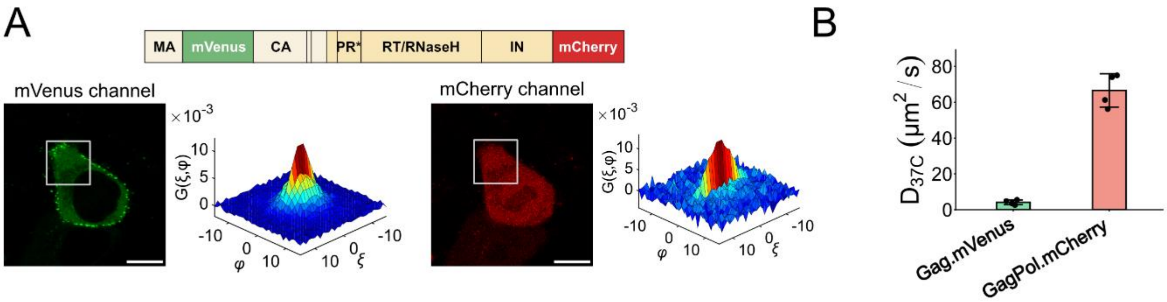

***Figure S3: Labeling GagPol at the C-terminal domain with mCherry. A)*** *Left: Representative image of HeLa cells transfected with the DL-IN-mCherry construct. The data were acquired using a laser power of 2 µW before the objective for both the mVenus and mCherry channels. RICS was performed by imaging the ROI highlighted by the white square and the SACF is reported on the right. Scale bar 10 µm. The schematic of the construct is shown above. Brightness and contrast are adjusted for visibility.* ***B)*** *Diffusion coefficients (mean ± standard deviation) of the GagPol DL-IN-mCherry construct shown in panel A. Results from 4 individual cells are shown as individual data points.*

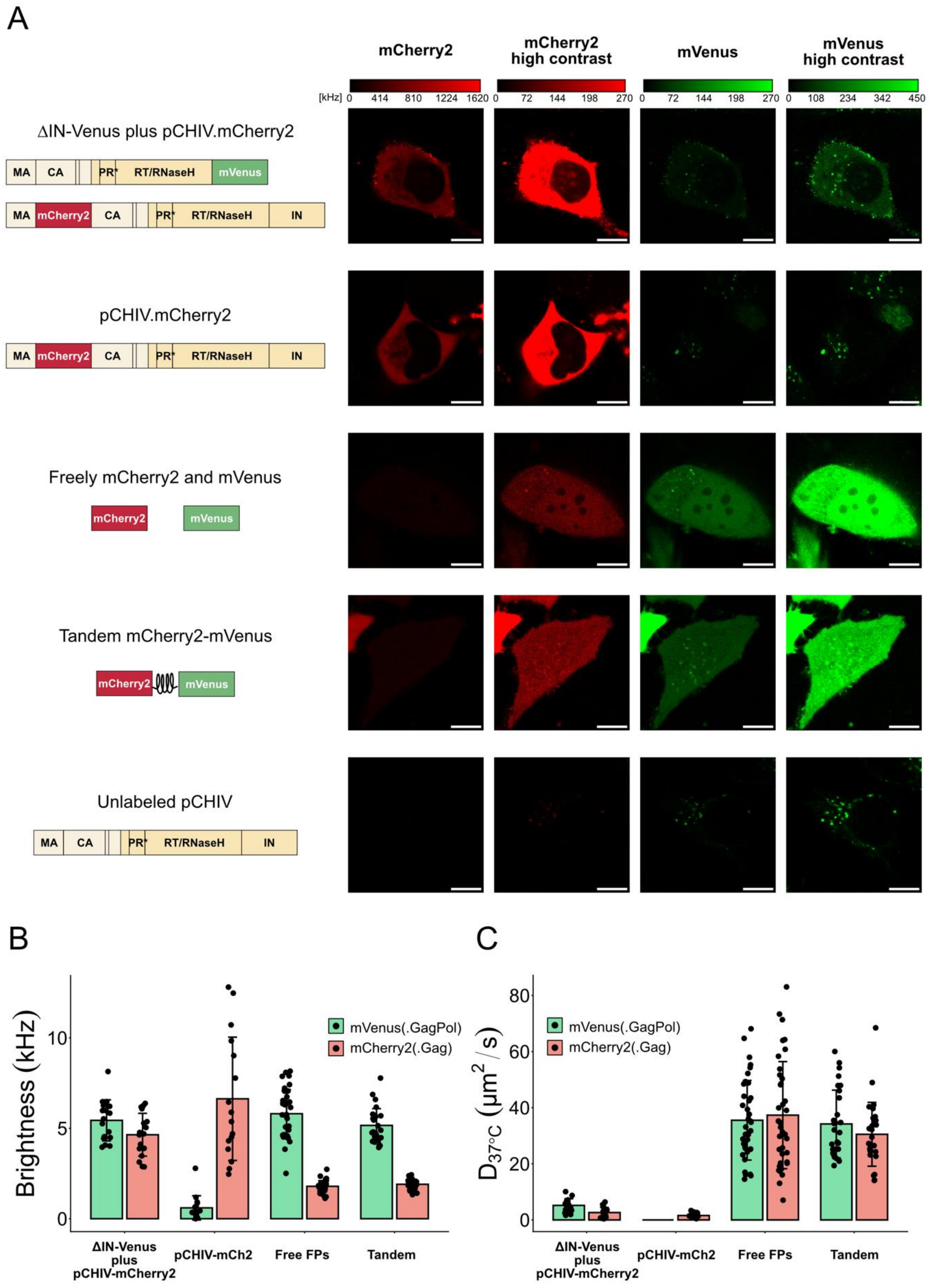

***Figure S4: Control RICS analyses – brightness and diffusion. A)*** *Representative images of HeLa cells expressing the construct depicted on the left and imaged in both the mCherry2 channel (595/50 nm emission) and mVenus channel (520/40 nm emission). Since all the acquisitions are done with the same parameters, but protein expression varies from cell to cell, images are reported with different contrast settings for visibility. The related contrast scale bar is added above the images. From top to bottom: ΔIN-Venus plus pCHIV-mCherry2 (same data shown in Figure 4A), pCHIV-mCherry2 (same data shown in SI figure 1), free mCherry2 and mVenus (mVenus data also shown in SI figure 1), mCherry2-mVenus tandem and unlabeled pCHIV as*

*control for autofluorescence.* ***B)*** *Molecular brightness (mean ± standard deviation) of the constructs depicted in panel A derived from the RICS analyses, as described in figure 4, and reported in Supplementary Tables S1 and S2.* ***C)*** *Diffusion coefficient (mean ± standard deviation) of the constructs depicted in panel A derived from the RICS analyses, as described in figure 4, and reported in Supplementary Tables S1 and S2.*

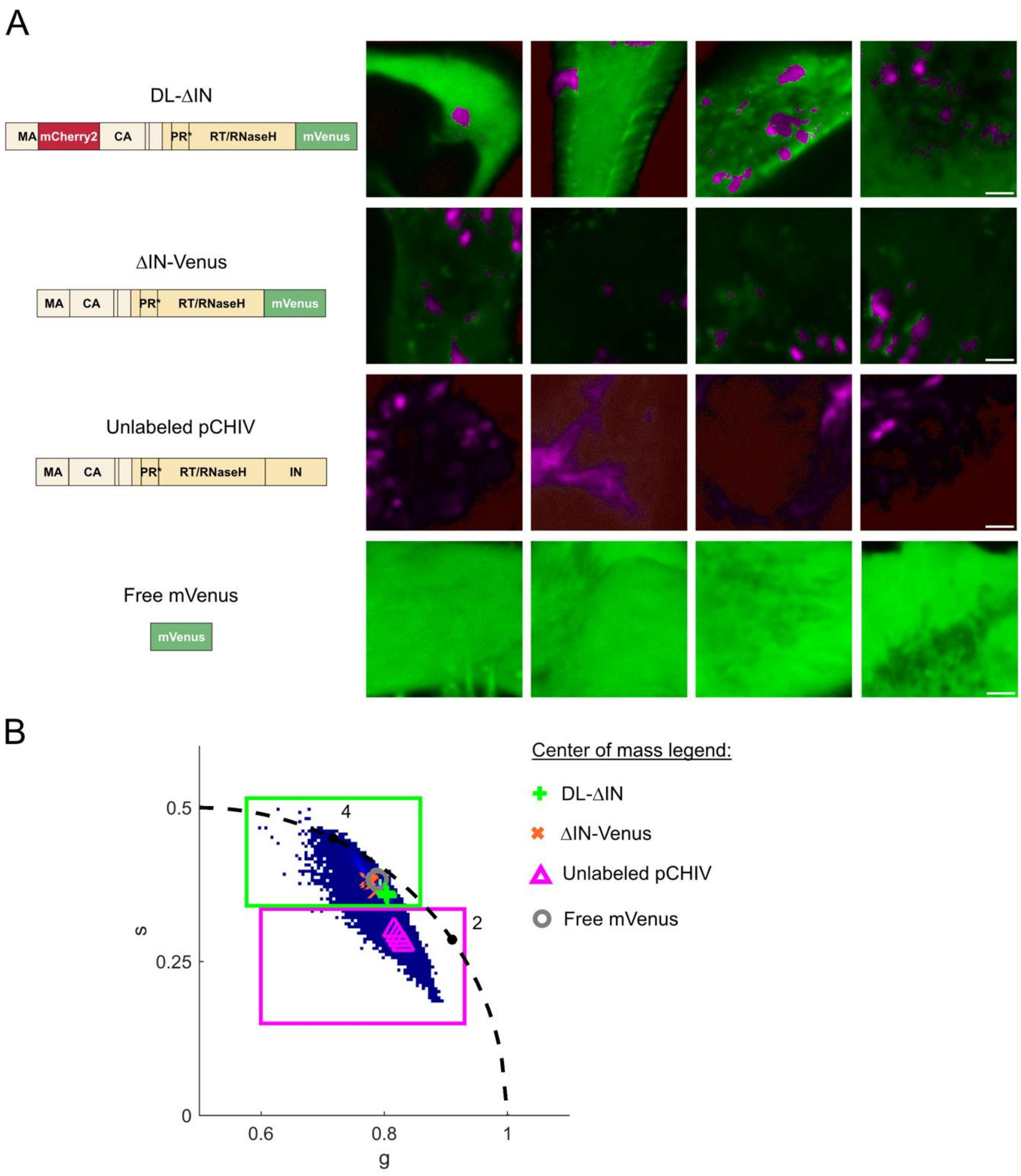

***Figure S5: FLIM analyses of GagPol.mVenus and cell autofluorescence. A)*** *Four individual HeLa cells transfected with various constructs. Scale bar 2 µm. Images are color-coded according to their fluorescence lifetime as defined in B, where magenta indicates the lifetime of autofluorescence component and green indicates the lifetime of mVenus. The contrast of each image was adjusted for readability.* ***B)*** *Phasor data from all individual files plotted together in a single phasor plot. The photon weighted center of mass of the phasor distribution for each individual file was calculated and plotted as symbols, allowing for direct comparison of the lifetime component across the datasets.*

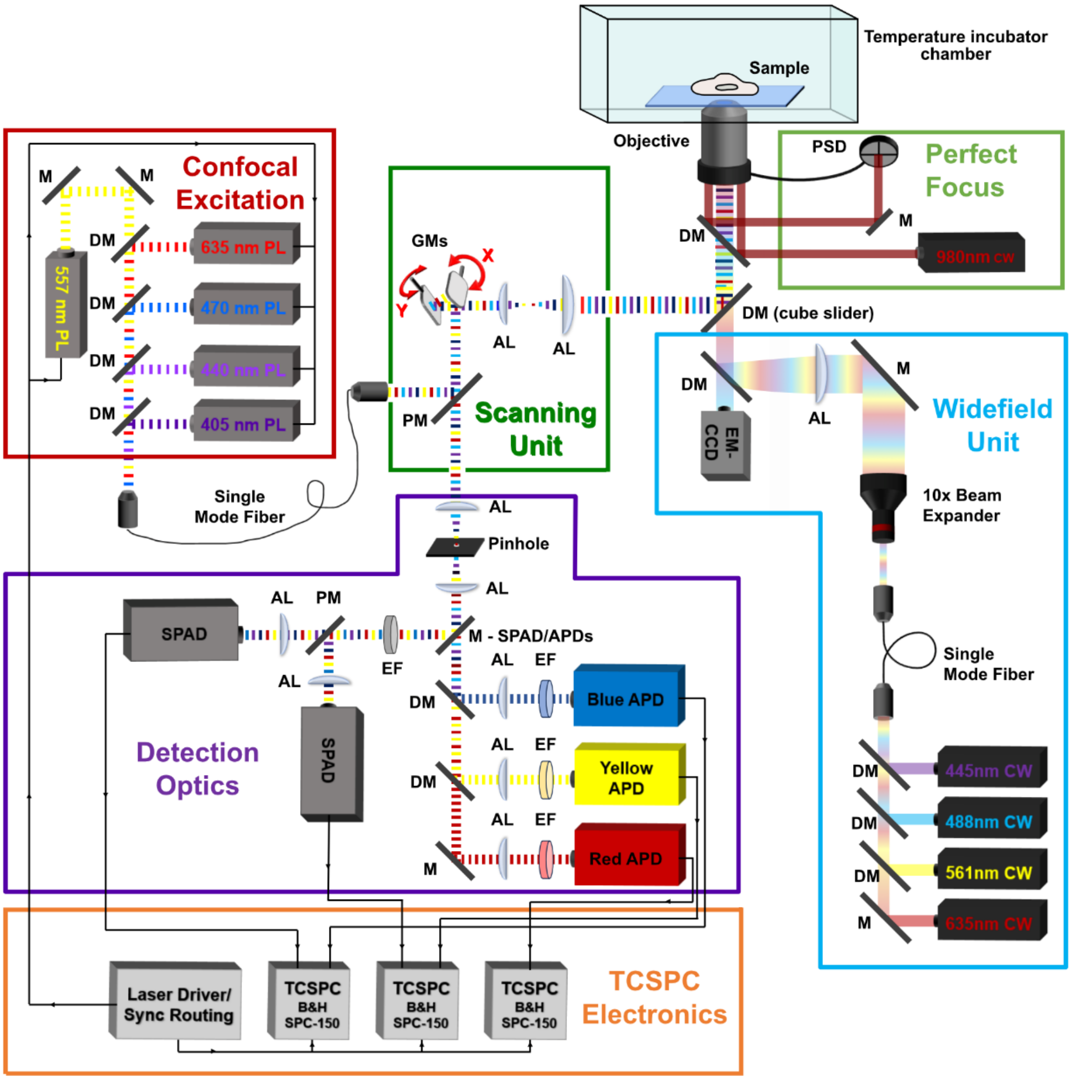

***Figure S6: Schematic illustration of the raster scanning confocal setup used in this work.*** *The microscope is equipped with five pulsed lasers (PL) of different wavelengths, driven by a picosecond pulsed driver (PDL 828 Sepia II, PicoQuant). The 470 nm and 560 nm PLs were used for confocal excitation of mVenus and mCherry2, respectively. The 440 nm PL was used for confocal excitation of PR-CFP-RT. All the pulsed lasers are coupled into a single mode fiber. At the fiber exit, the beam is collimated by a 20x apochromatic objective and reflected towards the scanning unit with a polychroic mirror (PM) (PM for 470/565 nm excitation: Semrock Di01-R405/488/561/635, AHF; PM for 445 nm excitation: Chroma zt442/514/561). Scanning is performed with galvanometric mirrors (GMs). After the GMs, the beam is expanded by a telescope and guided towards the objective (CFI Apo TIRF 100×, Nikon). The de-scanned emitted light is separated from the excitation light from the same PM, guided through an 80 µm pinhole towards the detector unit, composed of five detectors. Different wavelengths are separated via dichroic mirrors (DM) prior reaching the detectors. In this work, we used two avalanche photodiodes (APDs) preceded by a focusing achromatic lens (AL) and an emission filter (EF): 520/40 nm for mVenus detection (Blue APD) and 595/50 nm for mCherry2 detection (Yellow APD). The blue APD, with a 480/40 nm EF was used for CFP emission. The two Single Photon Avalanche Diodes (SPADs) detectors are accessible by inserting a mirror (M: SPAD/APDs) in the optical path. Excitation and detection are synchronized and controlled by time correlated single photon counting (TCSPC) electronics, composed of a TCSPC card per detector and the PDL laser driver. The microscope is also equipped with a widefield unit, used to simplify the localization of positively transfected cells. Continuous wave (CW) lasers are coupled into a single mode fiber and expanded 10-fold by two achromatic lenses (beam expander). Widefield fluorescence emission can be visualized with an EM-CCD camera or at the eyepiece. 488 nm and 561 nm CW lasers were used for widefield excitation of mVenus and mCherry2, respectively. Switching between widefield and confocal path is achieved by exchanging a dichroic mirror (DM) in a cube slider in the microscope body, without perturbing the alignment. The setup is equipped with a perfect focus system, achieved with a position sensing device (PSD) and a 980 nm laser. Live-cell imaging can be performed at 37 °C thanks to the incubator chamber.*

### Supplementary tables

**Table S1: Fit results of RICS SACF.mVenus of HeLa cells expressing the constructs of interest.**

| Construct | n | BR | $C_{10\text{-}90\%}$ µM | (D ± SD) µm²/s | (ε ± SD) kHz | $F_{mob}$ ± SD |
|---|---|---|---|---|---|---|
| PR-Venus-RT | 21 | 2 | 0.28-1.93 | 9.4 ± 2.5 | 7.3 ± 1.4 | 0.82 ± 0.08 |
| RT-Venus-IN | 19 | 2 | 0.32-1.25 | 8 ± 4 | 7.5 ± 2.0 | 0.83 ± 0.05 |
| DL-IN-mCherry (Gag.mVenus)[a] | 4 | 1 | - | 4.1 ± 1.3 | 4.6 ± 1.2 | 0.70 ± 0.14 |
| ΔIN-Venus (Pol.mVenus) | 17 | 3 | 0.04-0.14 | 5.2 ± 2.4 | 5.4 ± 1.1 | 0.75 ± 0.07 |
| DL- ΔIN (Pol.mVenus) | 29 | 6 | 0.19-0.57 | 2.8 ± 1.5 | 7.8 ± 1.6 | 0.68 ± 0.10 |
| PR-FL-Venus-RT | 20 | 2 | 0.27-1.45 | 8.4 ± 2.2 | 7.0 ± 0.7 | 0.84 ± 0.04 |
| PR-RL-Venus-RT | 13 | 2 | 0.40-1.26 | 10 ± 5 | 6.7 ± 1.3 | 0.84 ± 0.05 |
| PR-CFP-RT[b] | 28 | 3 | 1.09-4.98 | 7 ± 4 | 3.0 ± 0.6 | 0.64 ± 0.11 |
| mVenus | 37 | 4 | 0.36-2.44 | 36 ± 14 | 5.8 ± 1.3 | 0.94 ± 0.06 |
| mVenus within the tandem mCherry2-mVenus | 28 | 3 | 0.30-0.96 | 34 ± 12 | 5.2 ± 0.9 | 0.93 ± 0.05 |
| pCHIV.mCherry2[c] | 16 | 3 | - | - | 0.6 ± 0.7 | - |

Results from independent biological replicates (BR) are reported, with n indicating the overall number of cells analyzed. Concentration were calculated according to Equation 4, and are reported as the range of concentrations between the 10th and 90th percentile. The diffusion coefficient ($D$) and molecular brightness ($\varepsilon$) are calculated per each cell as described in the Materials and methods section, and the mean ± standard deviation (SD) is here reported. $F_{mob}$ represents the mobile fraction, calculated for individual cells and reported as mean ± SD. It should be noted that every SACF was fit with a two-component model (mobile and immobile) for consistency, even though several SACFs could be fit with just one mobile component.

[a] Contrary to the rest of the experiments, cells expressing DL-IN-mCherry were imaged with 470 nm laser at an intensity of 2 µW before the objective

[b] Contrary to the rest of the experiments, cells expressing the construct PR-CFP-RT were measured with 445 nm laser at an intensity of 2 µW before the objective.

[c] Control of autofluorescence, as cells expressing pCHIV.mCherry2 alone only present autofluorescence and cross-talk in the mVenus channel.

**Table S2: Fit results of RICS SACF.mCherry2 of HeLa cells expressing the constructs of interest.**

| Construct | n | BR | $C_{10\text{-}90\%}$ μM | (D ± SD) μm²/s | (ε ± SD) kHz | $F_{mob}$ ± SD |
|---|---|---|---|---|---|---|
| DL-IN-mCherry (Pol.mCherry)[a] | 4 | 1 | - | 67 ± 9 | 2.33 ± 0.20 | 0.93 ± 0.05 |
| DL- ΔIN (Gag.mCherry2) | 29 | 6 | 1.15-6.27 | 3.1 ± 2.7 | 6.6 ± 2.5 | 0.41 ± 0.16 |
| pCHIV.mCherry2[b] | 33 | 6 | 0.76-4.55 | 2.1 ± 1.5 | 5.6 ± 2.7 | 0.55 ± 0.16 |
| mCherry2 | 37 | 4 | 0.30-2.09 | 37 ± 19 | 1.8 ± 0.3 | 0.92 ± 0.08 |
| mCherry2 within the tandem mCherry2-mVenus | 28 | 3 | 0.37-0.93 | 31 ± 11 | 1.9 ± 0.3 | 0.90 ± 0.08 |

Results from independent biological replicates (NB) are reported, with n indicating the overall number of cells analyzed. Concentration were calculated according to Equation 4, and are reported as the range of concentrations between the 10th and 90th percentile. $D$ and $\varepsilon$ are calculated per each cell as described in the Materials and methods section, and the mean ± SD is reported. $F_{mob}$ represents the mobile fraction, calculated for individual cells and reported as mean ± SD. Every SACF was fit with a two-component model (mobile and immobile) for consistency, even though several SACFs could be fit with just one mobile component.

[a] Contrary to the rest of the experiments, cells expressing DL-IN-mCherry were imaged with 470 nm laser at an intensity of 2 μW before the objective

[b] In this table are reported the results of six biological replicates of HeLa cells expressing pCHIV.mCherry2. In three of these replicates, pCHIV.mCherry2 was co-expressed with ΔIN-Venus.

**Table S3: Fit results of ccRICS of HeLa cells expressing the constructs of interest and ratio of concentration corrected for brightness.**

| Construct | n | NR | ($RCA_{mVenus}$ ± SD) % | ($RCA_{mCherry2}$ ± SD) % | $N_{mVenus}/N_{mCherry2}$ | $N_{mCherry2}/N_{mVenus}$ |
|---|---|---|---|---|---|---|
| DL- ΔIN | 29 | 6 | 65 ± 17 | 8 ± 3 | 0.050 ± 0.022 | 23 ± 8 |
| ΔIN-Venus + pCHIV.mCherry2 | 17 | 3 | 35 ± 15 | 4.1 ± 1.7 | 0.047 ± 0.021 | 26 ± 12 |
| mVenus + mCherry2 (Free protein, negative control) | 37 | 4 | 7 ± 7 | 8 ± 6 | 1.6 ± 1.3 | 1.0 ± 0.8 |
| Tandem mCherry2-mVenus (positive control) | 28 | 3 | 51 ± 10 | 50 ± 7 | 0.85 ± 0.21 | 1.14 ± 0.24 |

Results from independent biological replicates (NR) are reported, with n indicating the overall number of cells analyzed. Relative Cross-correlation Amplitudes (RCAs) are calculated per each cell as described in the Materials and Methods section, and the mean value ± SD is reported. The ratio between mVenus-labeled proteins and mCherry2-labeled proteins was calculated after normalizing the $N$ value derived from the SACFs fit for the brightness of the monomeric fluorescent proteins (see Materials and Method for details). Here, the mean values ± SD are reported.

**Table S4: List of primers designed in this study and brief description of cloning strategies.**

| Primer name | sequence 5'-3' | PCR Template | Cloning strategy | Final construct |
|---|---|---|---|---|
| PRneg_D25N_1F | gcaattaaaggaagctctattaaacacaggagcagatg | Any pCHIV based construct | Generate PR negative pCHIV mutants using three fragments NEBuilder® HiFi DNA Assembly strategy. The D25N mutation is introduced from the sequence highlighted in red. | All PR negative pCHIV constructs generated in this study |
| PRneg_1R | gctgatacttctccttcactctcattgccactgtcttc | | | |
| PRneg_2F | cagaagacagtggcaatgagagtgaaggagaagtatcag | | | |
| PRneg_2R | ctacaccgaactgagatacctacagcgtgagcattgag | | | |
| PRneg_3F | ttctcaatgctcacgctgtaggtatctcagttcggtgtag | | | |
| PRneg_D25N_3R | atcatctgctcctgtgtttaatagagcttcctttaattgc | | | |
| BspEI-Venus_1392_Rev | taagcatccggagccagaccccttgtacagctcgtccatgccg | pmVenus | Addition of BspEI (Kpn2I) sites flanking mVenus. Kpn2I digestion of mVenus amplicon followed by ligation into digested construct PR-CFP-RT, containing CFP flanked by Kpn2I sites | RT-Venus-IN |
| BspEI-CFP-SGSG_Fwd | cccgctccggatctgggatggtgagcaagggcgaggagc | | | |
| BspEI-CFP-no-linker_Fwd | cccgctccggaatggtgagcaagggcgaggagc | pmVenus | Addition of BspEI (Kpn2I) sites flanking mVenus. Kpn2I digestion of mVenus amplicon followed by ligation into digested construct PR-CFP-RT. | PR-Venus-RT |
| BspEI-CFP-PR-site-Rev | cccgctccggaaggactaatgggaaaatttaaagtcttgtacagctcgtccatgccg | | | |
| BspEI-CFP-SGSG_Fwd | cccgctccggatctgggatggtgagcaagggcgaggagc | pmVenus | Addition of BspEI (Kpn2I) sites flanking mVenus and of a short flexible linker (SGSG) linker at the N-terminus. Kpn2I digestion of mVenus amplicon followed by ligation into digested construct PR-CFP-RT. | PR-FL-Venus-RT (FL : flexible linker consisting of one SGSG repeat) |
| BspEI-CFP-PR-site-Rev | cccgctccggaaggactaatgggaaaatttaaagtcttgtacagctcgtccatgccg | | | |
| BspEI-CFP_4P_Fwd | cccgctccggaccacctccacccatggtgagcaagggcgaggagc | pmVenus | Addition of BspEI (Kpn2I) sites flanking mVenus and of a short rigid linker (PPPP) linker at the N-terminus. Kpn2I digestion of mVenus amplicon followed by ligation into digested construct PR-CFP-RT. | PR-RL-Venus-RT (RL: rigid linker consisting of one PPPP repeat) |
| BspEI-CFP-PR-site-Rev | cccgctccggaaggactaatgggaaaatttaaagtcttgtacagctcgtccatgccg | | | |

| | | | | |
|---|---|---|---|---|
| mVenus_pCHIV_F | ggaaacaacagccagggatcgataggcggcatggtgagcaagggcgag | pmCherry2 | Amplification of mCherry2 sequence with the appropriate flanking sequences to be introduced between MA and CA of pCHIV via NEBuilder® HiFi DNA Assembly strategy. | pCHIV.mCherry2 |
| mVenus_pCHIV_R | gactatcgatccgcccttgtacagctcgtccatg | | | |
| BsiWI-mVenus_Fwd | agtgctggaatcaggaaagtactacgtacgatggtgagcaagggcgagga | pmVenus | Amplification of mVenus sequence with the appropriate flanking sequences to be introduced into pCHIV replacing the IN domain via NEBuilder® HiFi DNA Assembly strategy. | ΔIN-Venus |
| BsiWI-mVenus-Stop_Rev | cacaatcatcacctgccatctgttttccat*ttacgtacg*cttgtacagctcgtccatgcc | | | |
| GA_mVenus_19aa_fwd | cagtcgacggtaccgcgggcgtgagcaagggcgaggag | pmVenus | Amplification of mVenus sequence with the appropriate flanking sequences to be introduced into pmCherry2, also expanded by PCR, via NEBuilder® HiFi DNA Assembly strategy (see below) | Tandem mCherry2-mVenus |
| GA_mVenus_19aa_rev | ctagatccggtggatcccggcttgtacagctcgtccatg | | | |
| GA_mCherry2_19aa_fwd | ccgggatccaccggatctag | pmCherry2 | Amplification of pmCherry2 to be compatible with mVenus insert (see above) | |
| GA_mCherry2_19aa_rev | gcccgcggtaccgtcgac | | | |

All primers were purchased from Eurofin Genomics.
